# Composition and structure of synaptic ectosomes exporting antigen receptor linked to functional CD40 ligand from helper T-cells

**DOI:** 10.1101/600551

**Authors:** David G. Saliba, Pablo F. Céspedes-Donoso, Štefan Bálint, Ewoud B. Compeer, Salvatore Valvo, Kseniya Korobchevskaya, Viveka Mayya, Yanchun Peng, Tao Dong, Maria-Laura Tognoli, Eric O’Neill, Sarah Bonham, Roman Fischer, Benedikt M. Kessler, Michael L. Dustin

## Abstract

Cell communication through extracellular vesicles is an emerging topic in biology, including communication between cells of the immune system. Planar supported lipid bilayers (PSLBs) presenting T cell receptor (TCR) ligands and intercellular adhesion molecule-1 (ICAM-1) induce budding of extracellular microvesicles enriched in functional TCR, defined here as synaptic ectosomes (SE), from helper T cells. SE bind peptide-MHC directly exporting TCR into the synaptic cleft, but their ability to incorporate other effectors is unknown. Here, we utilized bead supported lipid bilayers (BSLB) to capture SE from single immunological synapses (IS), determined SE composition by immunofluorescence flow cytometry and enriched SE for proteomic analysis by particle sorting. Our results demonstrate selective enrichment of CD40 ligand (CD40L) and inducible T-cell costimulator (ICOS) in SE in response to addition of CD40 and ICOS ligand (ICOSL), respectively, to SLB presenting TCR ligands and ICAM-1. TCR triggering mobilized intracellular CD40L to the T cells surface at the IS, where it engaged CD40 to enable sorting into SE. SEs were enriched in tetraspanins and bone marrow stromal cell antigen 2 (BST-2) by immunofluorescence and TCR signalling and endosomal sorting complexes required for transport by proteomics. Super-resolution microscopy demonstrated that CD40L is present in microclusters within CD81 defined SE that are spatially segregated from TCR/ICOS/BST-2 microclusters. CD40L in SE retains the capacity to induce dendritic cell (DC) maturation and cytokine production. SE enabled helper T cells to release effectors physically linked to TCR.

**One Sentence Summary:** TCR and CD40L microclusters can be linked in synaptic ectosomes (extracellular vesicles) that are released in the immunological synapse by helper T cells and induce dendritic cell maturation and cytokine production.

## Introduction

Immune response communication depends on intercellular interactions of surface receptors expressed on T cells and antigen presenting cells (APC) *via* immunological synapses (IS), kinapses or stabilized microvilli (1, 2). In model IS, receptor-ligand pairs organize into radially symmetric supramolecular activation clusters (SMACs). The central (c)SMAC incorporates a secretory synaptic cleft, TCR interaction with peptide-major histocompatibility complex (pMHC) and costimulatory receptor-ligand interactions and is surrounded by the peripheral (p)SMAC enriched in LFA-1 (T cell side) interaction with ICAM-1 (APC side) enriched peripheral (p)SMAC (3). The dynamics of IS formation involves initial contacts through microvilli that trigger cytoplasmic Ca^2+^ elevation leading to rapid spreading and formation of SMACs through inward directed cytoskeletal transport (4, 5). Once the IS matures, TCR-pMHC pairs form in the distal (d)SMAC and segregate into microclusters (MCs) that integrate signaling as they centripetally migrate to the cSMAC where signaling is terminated (6). TCR MCs are a common feature of IS, kinapses and stabilized microvilli (1, 7). However, the IS is not only a platform for signal integration, but also enables polarized delivery of effector function. These include the polarized delivery of cytokines (8), nucleic acid containing exosomes (9), and TCR enriched extracellular vesicles that bud directly into the synaptic cleft from the T cell side of the IS (10). “Ectosomes” are extracellular vesicles released from the plasma membrane (11). Therefore, we define directly TCR elicited extracellular vesicles that are formed and simultaneously exported across the IS as synaptic ectosomes (SE).

CD40 ligand (CD40L, CD154) is a 39 kDa glycoprotein expressed by CD4^+^ T cells (12) and is one of the key effectors delivered by helper T cells through the IS (13, 14). Inducible T cell costimulator (ICOS, also known at CD278) interaction with ICOSL promotes CD40L-CD40 interactions in the IS (15, 16). CD40L is transferred to antigen presenting cells in vitro (17). Trimeric CD40L released by proteolysis by ADAM10 is a partial agonist of CD40, suggesting the fully active CD40 must remain membrane anchored to sufficiently crosslink CD40 for full agonist function (18, 19). How helper T cells achieve this high level of crosslinking in the IS is not established.

In this study we set out to determine the protein composition and mechanism of SE release in the synaptic cleft by helper T cells. To this aim we develop technologies for isolation of SE released by T cells directly at the IS on BSLB (20) and integrate complementary flow cytometry, mass spectrometry and super resolution microscopy data. We show that the polarized transfer of T cell derived SE is determined by selective sorting processes directly in the IS and depends on both the presence of ligands on the SLB and their segregation into the synaptic cleft, as shown for TCR complex:anti-CD3ε/pHLA-DR complexes, CD40L:CD40 and ICOS:ICOSL, but not LFA-1:ICAM-1 bound pairs. Other components, such as tetraspanins and BST-2, are enriched in SE without being engaged with a ligand. Quantitative mass spectrometry of SE revealed members of the core ESCRT machinery and adaptor proteins responsible for the scission of SE at the IS. Using direct stochastic optical reconstruction microscopy (dSTORM) we further demonstrate that SE contain discrete TCR/ICOS/BST-2 and CD40L microclusters. T cell budding of SE, therefore, provides a strategy to generate antigen specific and effector armed SE.

## Results

### CD40L is recruited to the IS and left by kinapses in a CD40 dependent manner

CD40L is stored in intracellular compartments within CD4^+^ effector cells and mobilized to IS where it engages CD40 in the IS upon activation through the TCR (21, 22). To mimic the APC surface and stimulate IS formation, the PSLB presented the adhesion molecule ICAM-1 and a Fab fragment of the anti-CD3ε mAb UCHT1 (UCHT1-Fab) (10), which functions like a strong agonist pMHC (23) (Figure 1A). Due to challenges with fluorescent protein tagging of CD40L, we detected it in the IS using an anti-CD40L mAb, which has the caveat that it competes with CD40, but nonetheless detects recruitment of CD40L to the IS (16). To determine the impact of CD40 density in the PSLB on detection of CD40L by this method we allowed IS to form on PSLBs presenting ICAM-1 and UCHT1-Fab over the physiological range of CD40 densities from 0 – 500 molec./μm^2^. The anti-CD40L signal was imaged by total internal reflection fluorescence microscopy (TIRFM) that only illuminates up to 200 nm into the sample, and thus restricts detection to the IS. Minimal IS CD40L was detected in the absence of CD40 as previously reported (16) and near uniformly increased anti-CD40L was detected at 10, 50 and 100 CD40 molec./μm^2^ with a reduction in signal at 500 CD40 molec./μm^2^ (Figure 1B). Thus, whether this loss of signal at high CD40 density is due to competition between CD40 and the anti-CD40L mAb or some other process, we conclude that CD40L can be detected and localized over the entire physiological range of CD40 densities using anti-CD40L antibody. To investigate the cellular localization of all CD40L, T cells were incubated on the PSLB with ICAM-1 and UCHT1-Fab without or with 50 CD40/μm^2^ for 30 minutes, fixed, permeabilized and stained with anti-CD40L (magenta) and CELL MASK^®^ (gray) to track cell membranes and 3D images generated by super-resolution Airyscan^®^ confocal microscopy. In the absence of CD40, CD40L puncta were polarized toward the IS, but were mostly 1-3 μm above the PSLB within the T cell (Figure 1C). In the presence of CD40, most of the CD40L was concentrated at the T cells-PSLB interface in a dense patch of bright puncta (Figure 1C). Live microscopy demonstrated that CD40L was detected in the IS just after and together with TCR-UCHT1-Fab complexes (Movie S1). It was not clear if these bright puncta of CD40L immunoreactivity were present in microclusters in the plasma membrane or if they were released into the synaptic cleft in some form. When T cells break IS symmetry and transition to migratory kinapses they leave trails of SE behind on the substrate (10) (Figure 1D). To investigate the fate of CD40L, we imaged IS and kinapses by TIRFM to avoid detection of intracellular CD40L. The cSMAC of IS contained partially overlapping signals for UCHT1-Fab and anti-CD40L staining as previously described (16) (Figure 1E; top panel; Movie S1 and S2). Upon kinapse formation by T cells, a trail of UCHT1-Fab was present without or with CD40 in the bilayer as expected (10) (Figure 1E; bottom panel, Movie S3). In the absence of CD40 in the PLSB, the few anti-CD40L reactive puncta that were detected remained associated with the migrating T cell (Figure 1F). Kinapses formed in the presence of CD40 resulted in a trail of anti-CD40L reactive puncta that remained tethered to the PSLB (Figure 1F). These images and quantification are consistent with CD40L release in SE along with or in parallel with the TCR.

**Figure 1:**
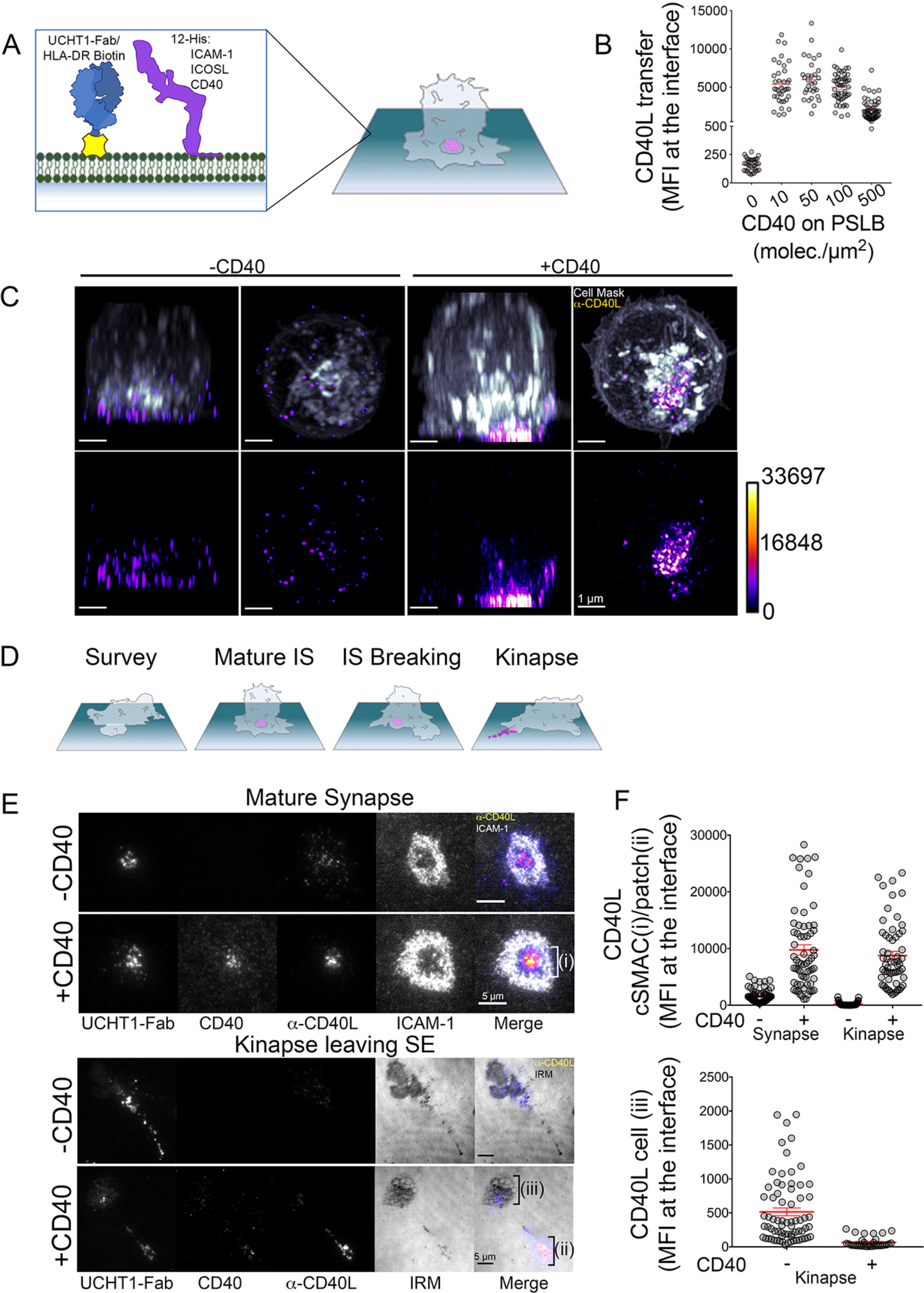
CD40 dependent recruitment of CD40L to the IS and deposition in SE trail. (A) Schematic of PSLB and mature IS. (B) Detection of CD40L with the anti-CD40L clone 24-31 as a function of CD40 in the PSLB. T cells were allowed to form IS for 10 minutes in the presence of Alexa Fluor^®^ 647 anti-CD40L antibody and imaged by TIRFM. Data is pooled from 5 donors with each point being one cell. (C) Representative Airyscan^®^ of CD4 T (Cell Mask, gray) cell releasing CD40L SE at the synaptic cleft on PSLB in the presence of CD40, Scale bar: 1μm. (D) IS and kinapse stages of T cell interaction. Stages of TCR positive SE are released at the synaptic cleft upon mature IS formation. Following symmetry breaking the SE are partly dragged by the kinapse as they are left (10). (E) Representative TIRFM of IS (top, 10 minutes incubation) and kinapse (bottom, 90 minute incubation) showing CD40 clustering in PSLB coated with ICAM-1, UCHT1-Fab in the presence or absence of CD40. Following fixation and permeabilization cells were stained with anti-CD40L, scale bar: 5 μm. (F) Detection of CD40L with anti-CD40L clone 24-31 in (D) (****p ≤ 0.0001) nonparametric Mann-Whitney test (U test). Data is from 5 donors.

### Selective transfer of CD40L and ICOS into SE

Uptake by APCs makes determining the composition of SE released by T cells challenging (10). In addition, the bidirectional transfer of membrane derived molecules between APCs and T cells engaged in the bipartite IS confounds the detection of loss of molecules from the T cell (10, 24). We developed a BSLB platform using the same compositions as on PSLB (Figure 2A) with similar synaptic accumulation of TCR as viewed by Airyscan^®^ confocal microscopy (Figure 2B vs Figure 1B, and Movie S4). Kinapse formation would not necessarily allow T cell to separate from BSLB, so we used low temperature and divalent cation chelation to inactive LFA-1-ICAM-1 interactions and gentle shearing forces provided by pipetting to separate the T cells from the BSLB. This enabled the quantitative profiling of gain of T cell molecules on BSLB and loss of molecules from the T cell surface by flow cytometry with a gating strategy based on light scattering and a fluorescent lipid signal to assure analysis of single BSLB (Figure 1 – figure supplement 1). As with the PSLB system, we needed to assess how CD40 on the BSLB would impact detection of CD40L in the IS using the anti-CD40L antibody. We evaluated this by flow cytometry over a range from 0-500 CD40 molec./μm^2^ (Figure 2C). We determined the relative IS transfer of CD40L (%) between T cells and BSLB using isotype control-corrected geometric mean fluorescence intensities (GMFI) as follows (GMFI_BSLB_ ÷ (GMFI_BSLB_ + GMFI_T cells_)) × 100 after 1.5 h incubation at 37°C and release of BSLB:T cell conjugates with ice cold PBS/EDTA. In the BSLB system, the decreased detection of CD40L using anti-CD40L mAb was not as prominent such that 500 CD40 molec/μm^2^ was used for most experiments (Figure 2C).

**Figure 2:**
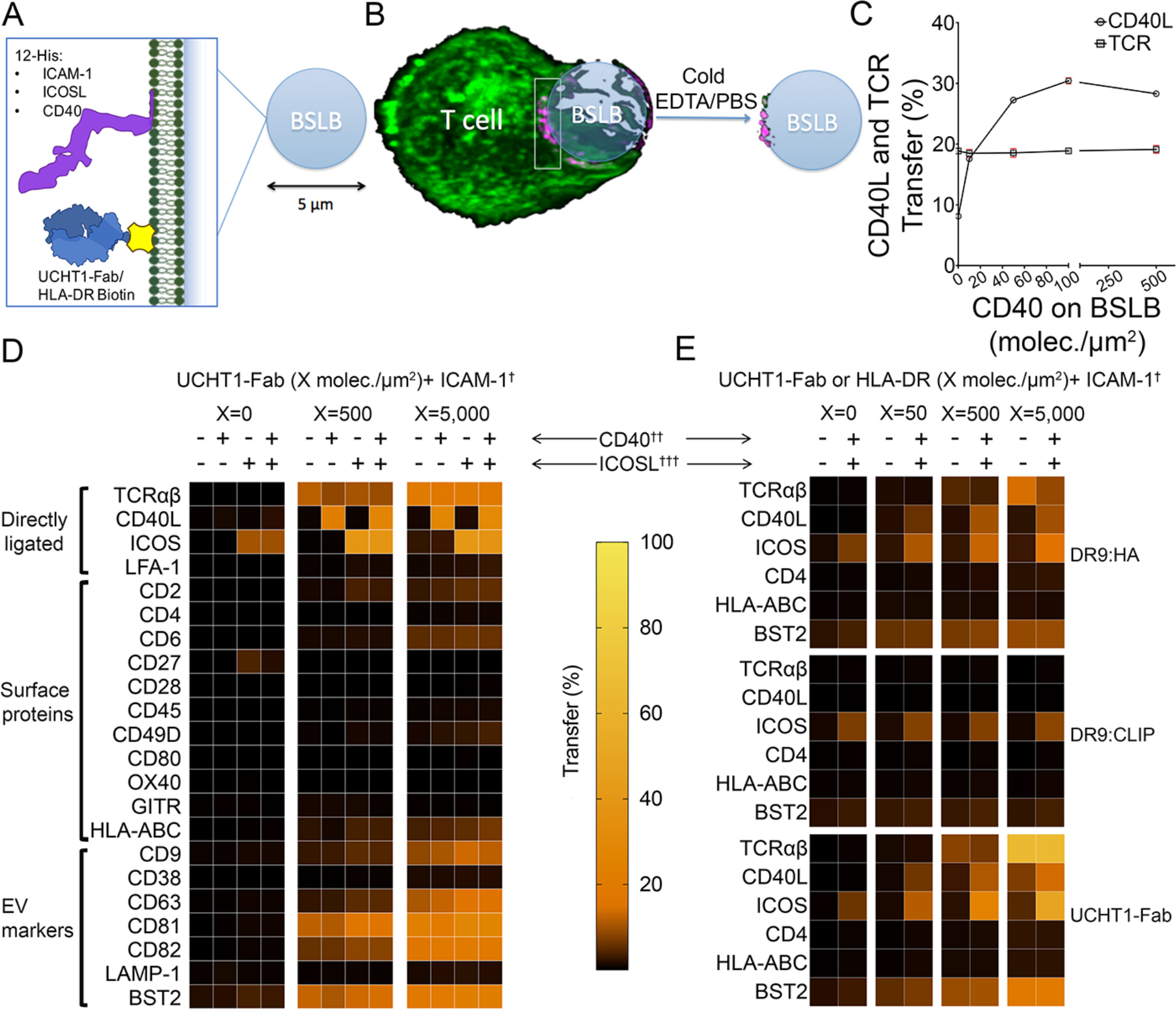
CD40L is incorporated in SE. (A) Schematic of BSLB (B) Schematic of T cell (green) interacting with BSLB with UCHT1-Fab (magenta) and SE on BSLB after T cell removal. (C) % CD40L and TCR transfer to BSLB coated with incremental levels of CD40. Data is from 6 donors. (D) Left-Heat maps showing the percentage of proteins transferred 5 from T cells to BSLB. Data is from 10 donors. (E)-Heat maps showing the percentage of proteins transferred from clone 35 to BSLB. Data is representative of 3 independent experiments with different clone 35 aliquots.

We next tested a larger panel of T cell surface molecules and calculated a relative IS transfer as above to assess selectivity of the sorting process leading to transfer of TCR, CD40L and other surface proteins by human CD4^+^ T cell blasts to BLSB presenting ICAM-1, UCHT1-Fab with or without combinations of CD40 and ICOSL representing an activated APC (Figure 2D, Figure 2 – figure supplement 2, Table S1). Transfer of TCR was ligand dose dependent and independent of ICOSL and CD40. ICOS was weakly transferred in response to ICOSL without UCHT1-Fab in the BSLB, but the efficiency was increased by TCR engagement. CD40L was transferred only when UCHT1-Fab and CD40 was present in the BSLB, and we were not able to detect a substantial increase when ICOSL was also included in the BSLB in contrast to prior results with PSLB (16) (Figure 2 – figure supplements 2 and 3). However, ICOSL presentation on BSLB increased conjugate formation between T cells and BSLB and also enhanced transfer of ICOS and tetraspanins (more significantly CD63) from T cell to BSLB in a manner depending upon the density of ICOSL and the dose of UCHT1-Fab (Figure 2 – figure supplements 2-4). We detected the enrichment of some additional proteins compared to the plasma membrane from which these vesicles bud. The most enriched (>10% transfer) class of non-ligated proteins are BST-2 (Tetherin) and the tetraspanins CD9, CD63, CD81 and CD82 (Figure 2D and Figure 2 – figure supplements 2 and 3). HLA-ABC was enriched in SE at an intermediated level (1-10%) along with other proteins including CD2, CD6 and CD49D). LFA-1, CD38 and LAMP-1 (CD107a) and CD4 were represented at <1% and were not enriched above the ordinary levels found in the plasma membrane (Figure 2D and Figure 2 – figure supplement 3D). To further test the specificity of enrichment we investigated the transfer of other T cell expressed proteins, including OX40, CD27 and GITR. None of these proteins were enriched on BSLBs, at least in the absence of ligands on the BSLB, demonstrating ligand-binding specificity as a dominant requirement for transfer. Its notable that LFA-1 is ligated by ICAM-1 on the BSLB and was not enriched in the SE from the CD4^+^ effector T cells (Figure 2D, left).

We next wanted to verify the location of proteins detected in SE transferred to BSLB in the IS formed on PSLB containing UCHT1-Fab, ICAM-1, ICOSL and CD40. The localization of BST-2, CD63, CD81, CD82 and CD40L all associated with the cSMAC (Figure 2 – figure supplement 5A). BST2 and the tetraspanin CD81 overlapped precisely with UCHT1-Fab, whereas the tetraspanins CD63 and CD82 partially overlapped (Figure 2 – figure supplement 5A). The UCHT1 and CD40L appeared to be systematically displaced from each other (Figure 2 – figure supplements 5A,B). Thus, all the SE associated signals from the BSLB transfer experiments localized in the cSMAC in the PSLB system as predicted, but co-localization was variable.

To extend our findings to the context of a physiological ligand for TCR, we investigated the dynamics of SE enrichment using an antigen specific helper T cell clone reactive to HLA-DRB1*09:01-influenza HA_338-355_ (Figure 2E and Figure 3). We used HLA-DRB1*09:01 loaded with CLIP peptide as a non-agonist MHC:peptide control, as well as UCHT1-Fab as a positive control. CD40L, TCR and BST2 were specifically transferred to BSLB coated with HLA-DRB1*09:01-influenza HA_338-355_ and UCHT1-Fab compared to HLA-DRB1*09:01:CLIP. On PSLB, CD40L localizes to the center of the IS predominantly in the presence of CD40 and HLA-DRB1*09:01-influenza HA_338-355_, not in the presence of HLA-DRB1*09:01:CLIP (Figure 3A,B). BST2 was also co-localized in the TCR in an antigen dependent manner (Figure 3C). As with the polyclonal CD4^+^ T cells, some ICOS transferred to BSLB with ICOSL with control HLA-DRB1*09:01:CLIP or in the absence of any MHC molecules (Figure 2E), a phenomenon which is accompanied by the recruitment of ICOSL to an TCR independent cSMAC-like structure on the PSLB (Figure 3C,D). This ICOSL driven TCR independent synapse may exert some control over migration of T cells, but it did not lead to CD40L transfer in any setting and thus does not appear to directly elicit delivery of T cell help.

**Figure 3.**
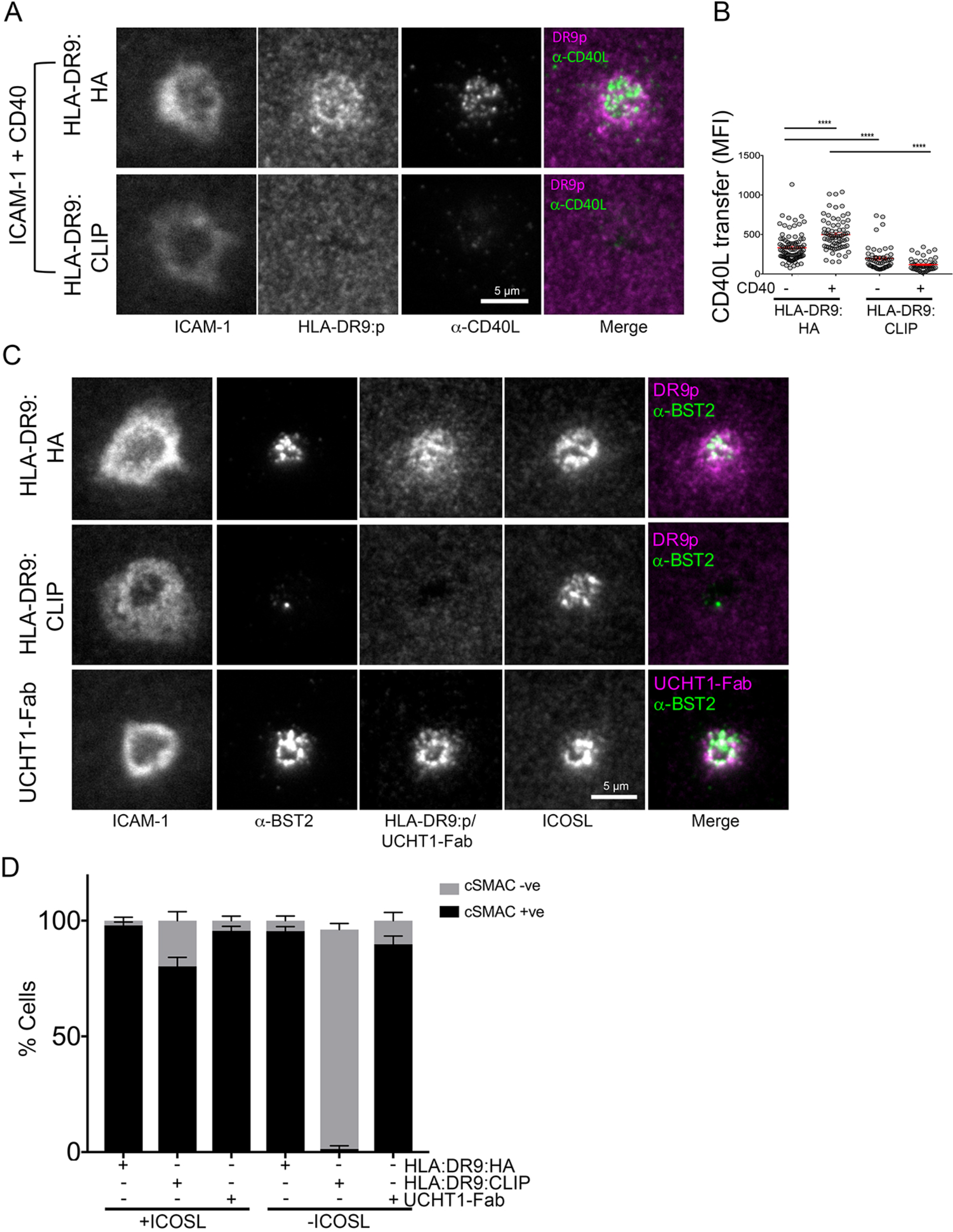
CD40L, BST2, and ICOSL localize to the synaptic cleft. (A) Representative TIRF images and IRM images showing staining of CD40L in the IS following incubation of HA specific clones on PSLB coated with ICAM-1 and either HLA-DR9_HA_ or HLA-DR9_CLIP_ monomers. Scale bar: 5μm. (B) Tukey box plot representation of CD40L staining from (A) expressed as arbitrary unit (A.U.). (****p ≤ 0.0001) nonparametric Mann-Whitney test (U test). Data is from 3 experiments with different clone 35 aliquots. (C) Representative TIRF and IRM images showing staining of BST2 in the IS following incubation of HA specific clones on PSLB coated with ICAM-1 and either His-tagged UCHT1-Fab, HLA-DR9_HA_ or HLA-DR9_CLIP_ monomers. Scale bar: 5μm. (D) Percentage cells with or without cSMAC like structure (as seen in C) following incubation of HA specific clones on PSLB coated with ICAM-1 and either His-tagged UCHT1-Fab, HLA-DR9_HA_ or HLA-DR9_CLIP_ monomers in the presence or absence of ICOSL.

Activated T cells have been shown to transfer CD40L to B cells that expressed CD40, but lacked cognate peptide-MHC in vitro (17). We thus wanted to ask if activated human T cell were also capable of transferring CD40L to BSLB that lack UCHT1-Fab, but present CD40. We prepared UCHT1-Fab presenting BSLB^Atto-488^ and UCHT1-Fab negative BSLB^Atto-565^ in the 4 possible combinations where each either presents or does not present CD40 (Figure 4A). TCR and CD40L were readily detected on the UCHT1-Fab and CD40 bearing BSLB at 1.5 h and 24 h (Figure 4A). The surface expression of CD40L on the T cell was detectable at 1.5 h and 24 h and was decreased when CD40 was also present on the BSLB with UCHT1-Fab. No CD40L was detected on BSLB when CD40 was not present (Figure 4A). This high degree of specificity suggests that CD40L transfer is tightly linked to IS formation, but we wanted to further investigate strongly general activation could trigger CD40L transfer to ICAM-1 and CD40 bearing SLB in the absence of TCR engagement. We followed two approaches. First, we incubated T cells with phorbol myristate acetate (PMA) and ionomycin for 30-min to expose CD40L on the surface and then for another 90 min in the presence of BSLBs with ICAM-1 and CD40 only or ICAM-1, UCHT1-Fab and CD40. PMA-ionomycin significantly increased the relative transfer of CD40L to BSLBs with 0 or 20/μm^2^ UCHT1-Fab (Figure 4B). This demonstrates that TCR engagement is not absolutely necessary for CD40L transfer. In this context, we also asked if CD40L transfer requires free ubiquitin, like TCR transfer into SE (10). We depleted free ubiquitin by pre-treatment with the proteasome inhibitor MG132 (50 μM for 0.5 hours). MG132 pretreatment reduced CD40L transfer significantly, but had only a marginal effect on TCR transfer (Figure 4B, C). Thus, we conclude that human CD4^+^ T cells restrict free ubiquitin dependent transfer of CD40L to antigen specific IS, but this can be overcome by strongly upregulating CD40L using a powerful pharmacological agonist.

**Figure 4.**
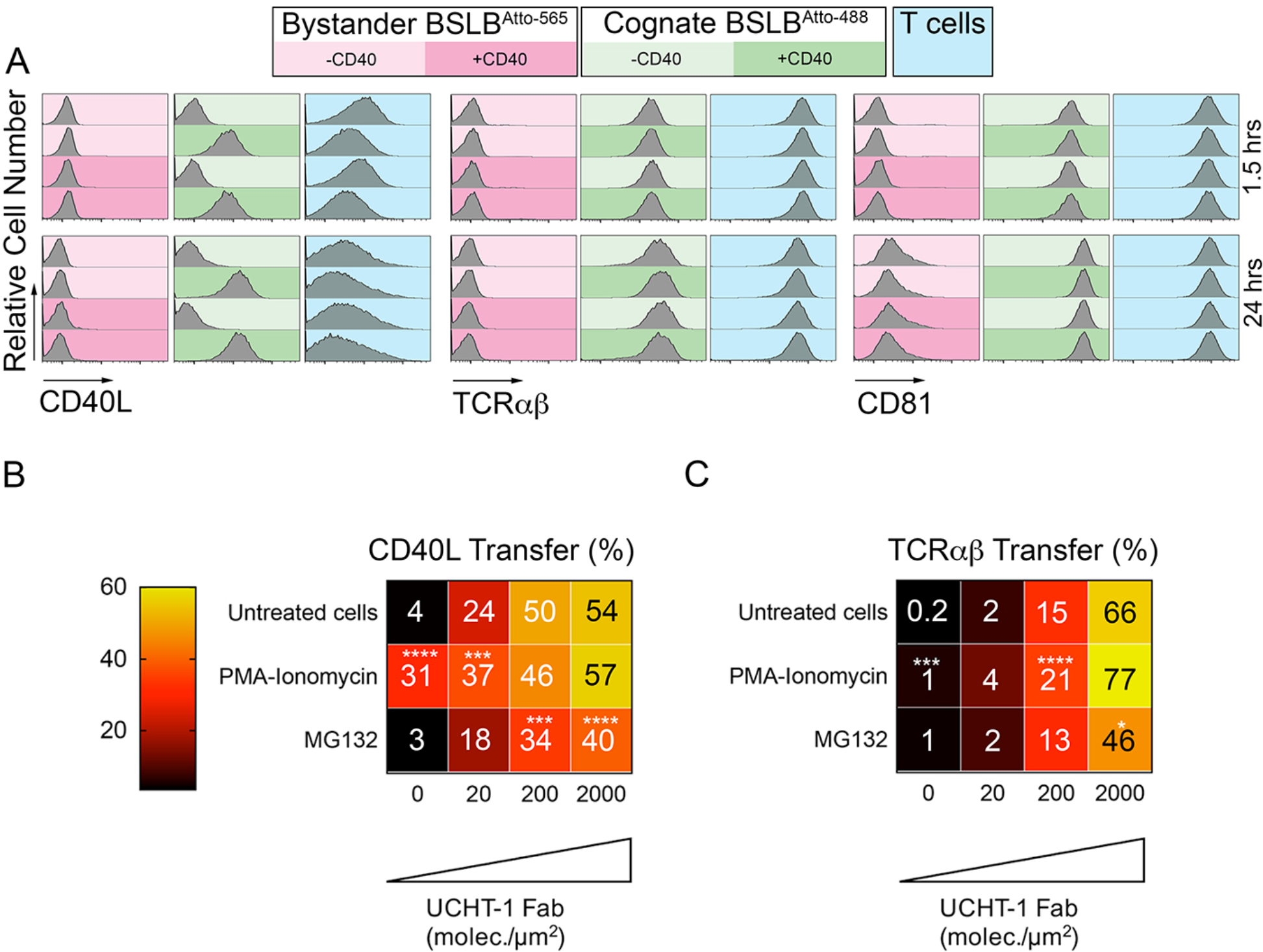
Specificity of CD40L transfer-effects of bystander BSLB, general activation and ubiquitin depletion. (A) Representative flow cytometry histograms and percentage marker transfer (CD40L, TCR and CD81) of “cognate” (UCHT1-Fab +ve Atto488; green) and bystander (UCHT1-Fab –ve Atto565; red) BSLB and T cells (blue). (B,C) Multiple t-test to compare the relative synaptic transfer (%) of CD40L (B) and TCRαβ (C) between cells pulsed for 30 minutes with either PMA-Ionomycin (10 ng/mL and 0.5 μg/mL) or MG132 (50 μM) and then incubated for another 90 min with agonistic BSLBs (increasing densities of UCHT-1-Fab, CD40 20 molec./μm^2^, ICAM-1 200 molec./μm^2^) in the presence of the drugs (values inside each cell represent mean percent synaptic transfer of 6 donors). Untreated cells were used as controls and as reference group for statistics; *, p ≤0.05; *** p ≤0.001; *** p ≤0.0001 (data is from 6 donors).

### Mass Spectrometry (MS) of SEs reveals enrichment of ESCRT proteins and TCR signaling

TCR-enriched SE are released through a TSG101 and VPS4-dependent plasma membrane budding process (10). Both TSG101 and VPS4 form part of the Endosomal Sorting Complex Required for transport (ESCRT). Specifically, TSG101 (an ESCRT-I member) is required for TCR sorting into membrane buds (6), whilst VPS4 mediates scission from the plasma membrane and release into the cSMAC/synaptic cleft (10). Residual ESCRT components may be trapped in the SE and the repertoire of ESCRT components involved in this process expanded by MS analysis of a sorted BSLB to enrich SE. We allowed 50 million CD4^+^ effector T cells to form IS on BSLB with ICAM-1, CD40, ICOSL in the presence or absence of UCHT1-Fab, disengaged the T cells and BSLB by incubation with ice-cold PBS/EDTA, sorted 5 million SE and subjected them to MS analysis after extraction of lipids. STRING analysis of 130 candidates having an absolute fold change (+UCHT1-Fab/-UCHT1-Fab) of at least 3.35 revealed a network of proteins with reported interactions. They are displayed to minimize the energy of the system based on the confidence score for interactions (Figure 5A, supplement excel file 1). A Markov clustering algorithm (25) highlighted six modules with 3 or more nodes displaying a high degree of adjacency (Figure 5B). We used Reactome (reactome.org) to determine if more proteins are found for particular pathways than is predicted by chance. This analysis revealed significant enrichment of TCR signaling, vesicle mediated transport and ESCRT machinery pathways (Figure 5C). Several surface proteins identified as enriched in the immunofluorescence analysis (Figure 2D) were also enriched in SE as defined by MS including CD40L, ICOS, CD81, CD82, BST2, HLA-A, HLA-B, CD2, CD6 and CD49d (ITGA4/ITGB1) (Figure 5A, and supplement excel file 1). Most of the surface proteins that were not enriched in the immunofluorescence analysis (Figure 2D) were also not detected in the MS analysis including CD4, CD27, CD38, CD45, CD80, OX40, GITR and LAMP1 (Figure 5A, and supplement excel file 1). However, a few proteins not enriched in SE by immunofluorescence analysis (Figure 2D) including LFA-1 (ITGAL/ITGB2) and CD28 were enriched in SE by MS (Figure 5A, and supplement excel file 1). However, reexamination of the raw immunofluorescence data reveals that both LFA-1 and CD28 gave a stronger signal on BSLB with UCHT1-Fab compared to without UCHT1-Fab, even though only 0.2% of the surface protein was transferred in the presence of UCHT1. Thus, the MS analysis fully agreed with the immunofluorescence analysis. ESCRT-0 components HRS, STAM2 and EPN1 and ESCRT-II component ALIX were enhanced on BSLB by TCR engagement (Figure 5A, and supplement excel file 1). Within the range of TIRF microscopy, EPN1 and ALIX were localized in or around the cSMAC (Figure 5D,E), whereas STAM2 was present in the cSMAC, but also in peripheral components of the IS, consistent with additional functions of STAM2 in the IS (26). MS also revealed enrichment of disintegrin and metalloproteinase domain-containing protein 10 (ADAM10), which mediates CD40L release as a soluble trimer from T cells upon CD40 engagement (18), on SE. The BSLB system thus expands our understanding of SE composition, including candidates for ESCRT pathway function and other aspects of SE biology.

**Figure 5.**
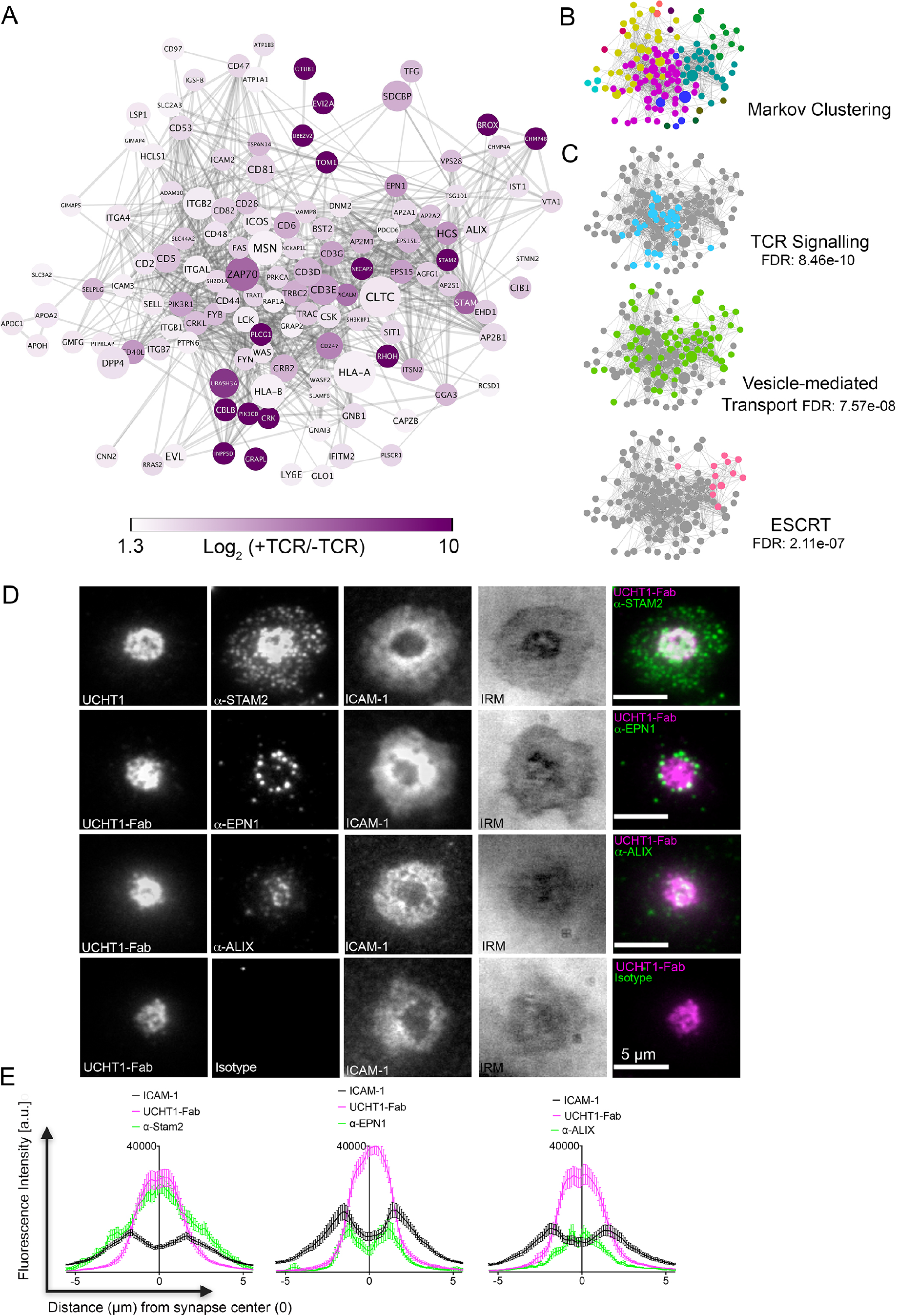
SE contains ESCRTs and TCR signalosome. (A) Proteins enriched by UCHT1-Fab on BSLB also containing ICAM-1, ICOSL and CD40. The network plot is based on known and predicted interactions from the STRING database (v11), with minimal confidence score of 0.4. Each protein displayed >1.75 log2 fold enrichment over BSLB coated with ICAM-1, ICOSL and CD40 in two independent experiments (3 pooled donors/experiment). Node area represents protein LFQ intensity. Line thickness represents confidence score (0.4-0.999). False discovery rate (FDR) on peptide and protein level were set to 1%. (B) Modules identified in protein network shown in (A) determined using the Markov Cluster algorithm (inflation parameter: 2.5). Each colour represents a separate module (associated adjacency matrix). (C) Reactome analysis (reactome.org) of protein network shown in (A) reveals enrichment of TCR signalling, Vesicle-mediated transport and ESCRT pathways. (D) Representative TIRFM and IRM images showing staining of STAM2, EPN1 and ALIX on PSLB with UCHT1-Fab and ICAM-1, CD40 and ICOSL. T cells were incubated with PSLB at 37°C for 1 hour. Following fixation and permeabilization, cells were stained with relevant antibodies. (E) Average radial distribution in IS formed on PLSB containing ICAM-1 (black), ICOSL, CD40 and UCHT1-Fab (magenta) stained with antibodies to the candidate proteins (green). Data is from 3 donors and 50 cells; scale bar: 5μm

### TCR and CD40L occupy spatially distinct microclusters within single SE

We next investigated the nanoscale organization of protein microclusters in SE deposited on PSLB after gentle removal of helper T cells (Figure 6A). We applied three-color dSTORM to localize SE proteins with a resolution of 20 nm. To visualize SE we took advantage of the fact that these membranous structures are enriched in glycoproteins compared to the sparsely populated PSLB and used wheat germ agglutinin (WGA) labelling to provide contrast. In agreement with earlier electron microscopy measurements (10), the average diameter of SE was 84 ± 5 nm (Figure 6B-D) and 36 ± 3 SE were transferred per IS (Figure 6E). The number and size of TCR microclusters transferred on SEs is similar to that of TCR microclusters present in the early IS (Figure 6F-H). This provides further evidence that TCR microclusters are converted into SE. To determine the localization of protein microclusters the SE were stained with anti-CD81-CF568 (Figure 7A), anti-TCRαβ-AF488 and a mAb for BST2, CD40L, or ICOS conjugated with AF647. Three color dSTORM revealed 3 subsets of SE: 1) TCR only, 2) CD40L, ICOS or BST2 only and 3) double positive for TCR and CD40L (54.5%), ICOS (77.5%) or BST2 (75.0%) (Figure 7B and Figure 7 – figure supplement 1-3). To address the degree of colocalization we applied two independent methods: coordinate-based colocalization (CBC) (27) and cross-correlation (28). TCR clusters were colocalized with ICOS or BST2 within a search radius of 50 nm, and segregated from CD40L clusters (Figure 7C). There was a high correlation between TCR with ICOS or BST2 at inter-particle distances less than 50 nm, whereas TCR and CD40L showed the highest correlation at a distance of 150 nm (Figure 7D and Figure 7 – figure supplement S3). Corroborating this, the mean nearest-neighbor distance (NND) of paired single-molecule localizations was 34 nm for TCR and BST2, 17 nm for TCR and ICOS and 230 nm for TCR and CD40L (Figure 7E). As a positive control, TCR molecules on SE detected with anti-TCRαβ−AF488 and UCHT1-Fab-AF647 showed a high degree of colocalization (Figure 7C-E). The results are consistent with TCR, BST2 and ICOS occupying overlapping microclusters, whereas TCR and CD40L occupied spatially distinct microclusters that are sorted within single SE. Segregation of microclusters within SE and populations of single positive vesicles explain the failure of TCR and CD40L immunofluorescence to co-localise in IS images (Figure 2-figure supplement 5 and Figure 3), although they were otherwise closely linked in the cSMAC.

**Figure 6.**
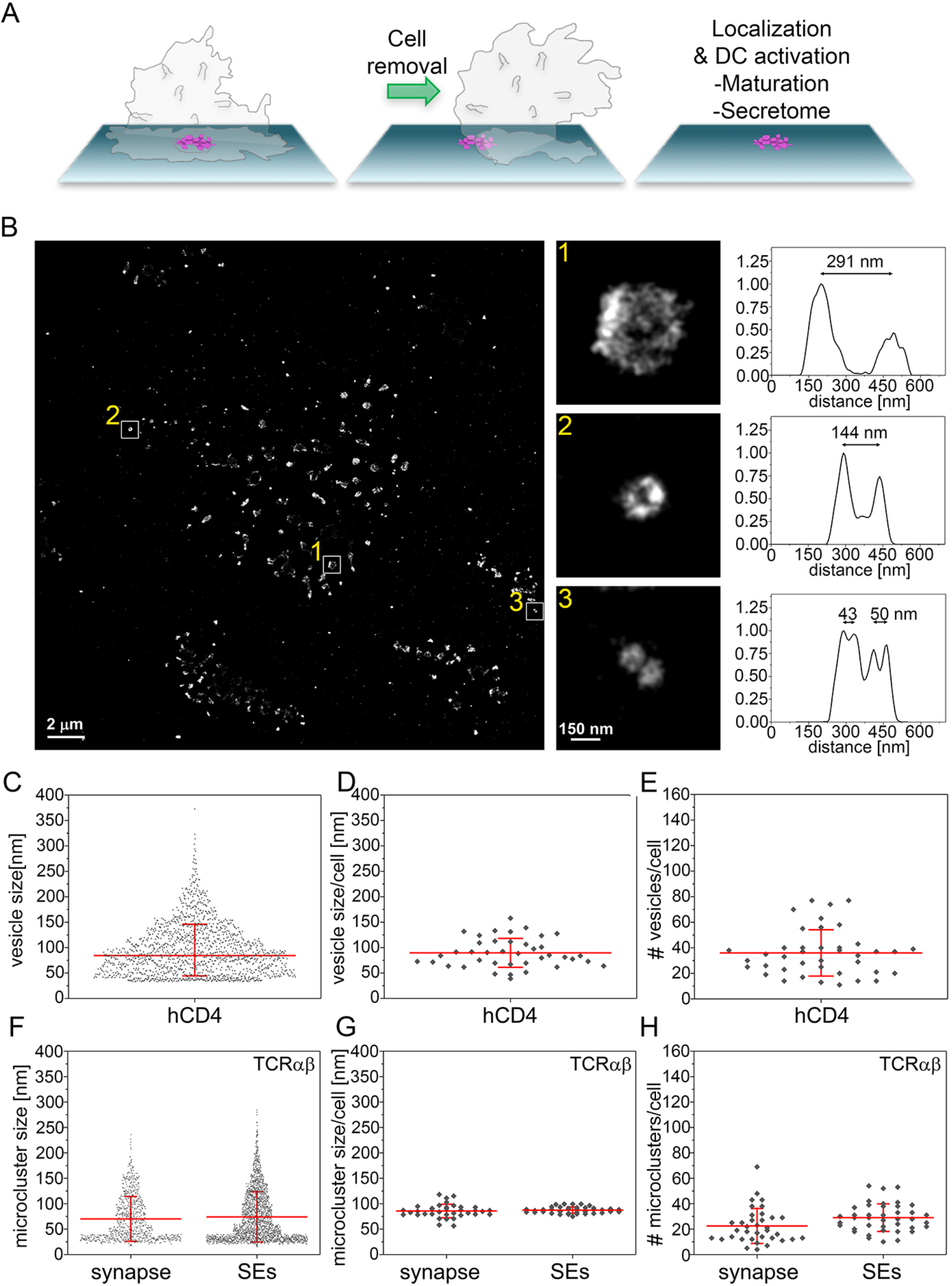
Size distribution SE captured from immunological synapses on PSLB. (A) Schematic of SE deposition on PSLB. (B) Representative dSTORM image of SE released by CD4^+^ T cells incubated for 90 min on supported lipid bilayers coated with ICAM-1, UCHT1-Fab, CD40 and ICOSL. SE were stained with WGA-CF568 to visualize the lipid membrane. Examples of SE of different sizes, depicted by the white squares, are zoomed-in and shown with correspondent relative fluorescence intensity profiles. (C) Size distribution of SE released from all cells. Each symbol represents an SE (n = 1482). (D) Size distribution of SE released per cell. Each symbol represents the median size of SE released per cell (n = 40). (E) Number of SE released per cell. Each symbol represents the median number of SE released per cell. (F) Size distribution of TCRαβ microclusters from synapse and SE. Each symbol represents a microcluster. (G) Size distribution of TCRαβ microclusters from synapse and SE released per cell. Each symbol represents the mean size of SE released per cell. (H) Number of TCRαβ microclusters from synapse and SE released per cell. Each symbol represents the mean number of microclusters per cell. Lines and errors represent means ± SD.

**Figure 7:**
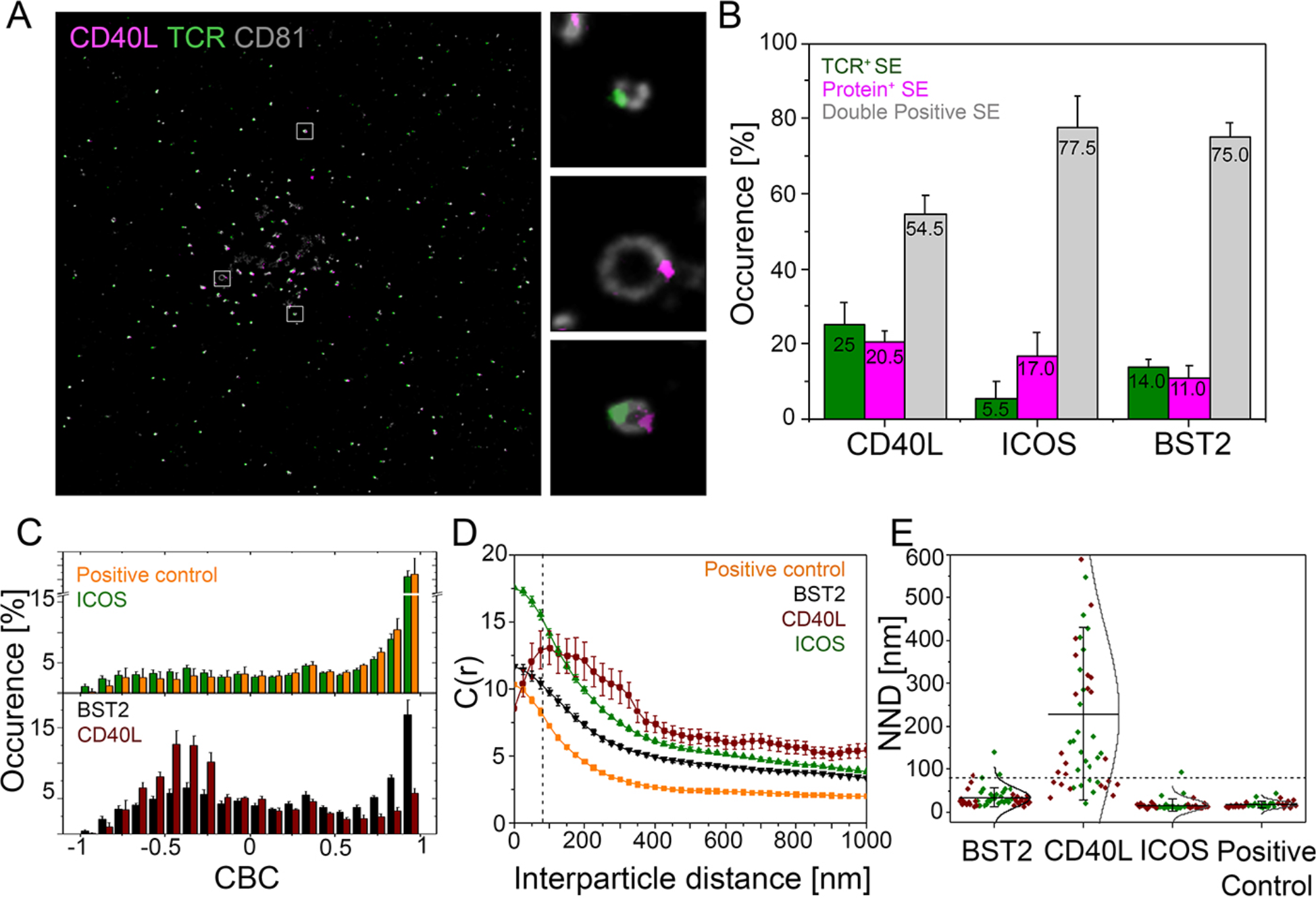
Nanoscale structure of SE. (A) Representative dSTORM images showing TCR (green), CD40L (magenta) on CD81 (gray) labeled SE. Insets show examples of SEs containing only TCR, only CD40L or both proteins. (B) Percentage of SEs containing only TCR (green), only protein of interest (magenta), or containing both TCR and protein of interest (grey). (C) CBC histograms of the single-molecule distributions of the colocalization parameter. Bars represent means ± SD. (D) Cross-correlation analysis. (E) Nearest-neighbor distance (NND) analysis from data shown in B). Each symbol represents the median NND of all paired single-molecule localizations from EVs released per cell. Lines and errors represent means ± SD. Dashed line in panel (D) and (E) marks mean size of SE.

### Released SE induce DC maturation and cytokine production

We next evaluated whether T cell isolated SE have an activating effect on monocyte-derived dendritic cells (moDCs). We therefore, first generated T cell derived SE on PSLBs coated with a combination of different T cell accessory ligands. After removal of T cells, immature moDCs were incubated with the released SEs for 24 hours followed by analysis of DC maturation markers and secreted factors. Conditions A1 and C0 had no SE, B1 had CD40L low SE, and C1 had CD40L high SE. CD40L high SE triggered high expression levels of HLA-DR and CD83 (Figure 8A, B and Figure 8 – figure supplement 1 A, B). Analysis of 105 secreted factors from DC, revealed that CD40L high SE significantly increased release of 46 factors, prominently including several chemokines and inflammatory cytokines such as tumor necrosis factor (TNF) and IL-12p70, which are important pro-inflammatory cytokines from licensed DCs (29) (Figure 8C and Figure 8 – figure supplement 1C). These results demonstrate that SE have an active form of CD40L that is sufficient to induce DC maturation in the absence of T cells. Another important aspect of SE mediated transfer is that many receptor systems involved in cell-cell communication requires a high order clustering, and therefore valency, to achieve full agonist function (19). The higher density of proteins in SE may therefore provide mechanistic insight in which SE promote extensive CD40 cross-linking on DC surface to robustly trigger down-stream signaling. Since each vesicle has an average diameter of 84 nm and there is an average of 37 vesicles released per IS (Figures 7C and D), the released surface area of all SE can be estimated as 0.82 μm^2^. This is ~0.2% of the surface area of the T cell, consistent with the % transfer of CD4 and CD38. Given this small surface area, the density of CD40L on SE is estimated to be over 200 trimers/μm^2^ (Table S3). As we have noted in the dSTORM imaging, these free CD40L trimers are further organized in even higher density microclusters. This provides a mechanism for SE to promote extensive CD40 cross-linking on the APCs surface to robustly activate down-stream signals. To test this hypothesis, we developed synthetic unilamellar vesicles (SUVs) loaded with increasing quantities of N-terminal His-tagged hCD40L (>200 molec./μm^2^, Figure 8D, Tables S3 and S4) to corroborate whether high binding valency due to CD40L concentration on sub-100 nm vesicles has a superior agonist effect on APCs. We then compared moDC activation when presented with either i) SUVs loaded with CD40L (vesicular CD40L), ii) soluble CD40L trimers (sCD40L) or iii) soluble CD40L trimers in the presence of unloaded (mock) SUVs. We observed a significant upregulation of HLA-DR, CD40, ICAM-1, CD80 and CD86 on moDC treated with vesicular CD40L compared to an equivalent concentration of soluble CD40L trimers (Figures 8E-F) demonstrating that vesicular CD40L is a superior agonist compared to soluble CD40L.

**Figure 8:**
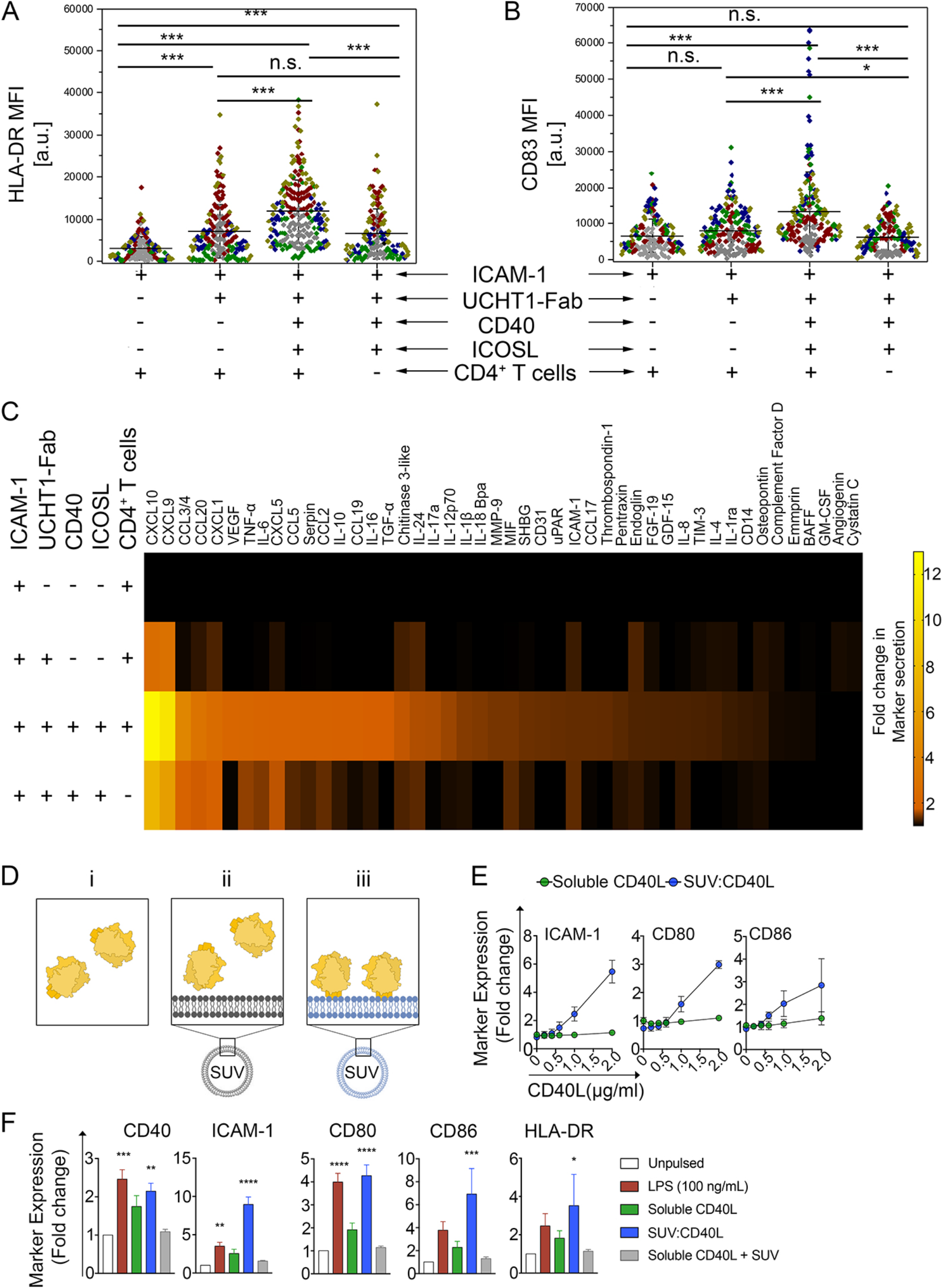
CD40L-positive SE left by T cells help DC and high-density vesicular CD40L is sufficient for DC maturation. (A, B) HLA-DR (A) or CD83 (B) expression on the surface of DCs stimulated for 24 h on PSLB prepared as indicated to present SE. Each symbol represents the mean fluorescent intensity of a cell (n ≥ 20) from 5 independent donors. Lines and errors represent means ± SD. ns, not significant; *, p ≤ 0.05; ***, p < 0.001; Kruskal-Wallis with Tukey`s post-hoc test. (C)After 24 h of culture, the supernatant of DCs stimulated as in A) and B) were analyzed for the presence of different secreted factors. Results from 7 independent donors. The fold-change was normalized to the results from the supernatant of DCs incubated with PSLB containing ICAM-1 alone. (D) For assessing the biological significance of presenting CD40L in vesicular structures compared to the proposed biologically active form of soluble trimeric CD40L [i, green], we developed 86 nm diameter synthetic unilamellar vesicles (SUV; Table S4) using phospholipids either without (ii, grey) or with His-tag, and hence CD40L, binding activity (iii, blue). An equal total mass (Mass eq.) of N-terminal His-tagged recombinant human CD40L was either incubated with DOPC liposomes (mock-SUV [ii, grey]) or used to load 12.5% DOGS-NTA and make vesicular CD40L (SUV:CD40L [iii, green]). As control, a Mass eq. of CD40L was used as soluble, free protein in culture. (E) FCM analyses of moDCs stimulated for 24h with either SUVs containing increasing amounts of CD40L monomers (blue; approximately 0, 40, 80, 120, 190 and 230 CD40L molec./SUV respectively) or an equivalent concentration of soluble CD40L (green). Data represent fold change in the GMFIs for ICAM-1, CD80 and CD86 of treated moDCs over unpulsed controls. (F) As in B, moDCs were stimulated using SUV:CD40L at a final load of approximately 230 molec./SUV. An equivalent concentration of soluble CD40L either delivered alone or in combination with SUVs lacking NTA lipids (soluble CD40L + SUV) were used as controls. After 24h of stimulation moDCs were collected, FcR blocked, stained and analysed by multicolor FCM for the expression of maturation markers. Given the high variability of arbitrary fluorescence values, the response was normalized as a fold change compared to unpulsed, immature DC controls. Data represent mean +/-SEM collected from 5 different donors and 5 independent experiments. One-way ANOVA and Dunn’s multiple comparisons test to the mock treated control was performed. *P* values < 0.05 (*); <0.002 (**); <0.0002 (***); <0.0001 (****) were considered significant.

## Discussion

In this study, we systematically profiled composition, nanoscale structure and function of trans-synaptic SE that were produced and captured in a model IS. Importantly, we adopted BSLB as a scalable platform to analyze SE by flow cytometry and mass spectroscopy. BSLB have been used previously to demonstrate activity of purified MHC proteins (30), detect weak protein interactions (20) and determine biophysical requirements for phagocytosis (31). We have also used BSLB to calibrate site densities of recombinant proteins attached to SLB by flow cytometry (32) and for bulk functional assays (6). Here, we allowed T cells to form IS with BSLB at a 1:1 ratio and then disrupted the conjugates to allow analysis of material transferred from the T cell to the BSLB. The strength of this approach is that each BSLB carries the output from one T cell, on average, such that the output of single IS can be quantified and calibrated to number of antibody binding sites and compared to average levels of the same molecules on the donor T cells. Another strength is that BSLB can then be isolated by particle sorting, which enabled MS analyses with minimal cellular contamination. A limitation of the approach is that while we can readily analyze both the cells and BSLB at a population level, we cannot link the cellular donor to a BSLB recipient specifically, which would require a different approach-perhaps with microfluidics. For these studies we used ~5 μm beads, which are near cell size and are easily analyzed on commercial flow cytometers. The results we obtained are qualitatively comparable to results with PSLB, but we cannot rule out some impact of the surface curvature of the BSLB. We also had some concern that the methods used to disrupt conjugates (divalent cation chelation and low temperature) might result in physical tearing off of membranes that might not normally be transferred to APC, and would contaminate SE. However, there is a high degree of enrichment for ligated cargo like ICOS and CD40L (>10%), tetraspannins (33) and BST2 (34) (~10%), and proteins with known tetraspannin association, like MHC class I (1-10%). In contrast, most other proteins are transferred at a level similar to the area of membrane transferred (~0.2%). This suggests a high degree of selectivity of the transfer process and minimal contamination from random pieces of plasma membrane. We previously had failed to detect tetraspanin CD63 enrichment in SE (10), but find here tetraspanin CD81 in the cSMAC, and enrichment and uniform distribution in TCR and/or CD40L positive SE by dSTORM. BST2 may tether some SE to the T cell membrane and account for dragging of SE behind kinapses, which also has implications for signaling of SE induced signaling to the T cell (34, 35). We observed formation of TCR engagement independent IS-like structures and SE-like vesicle transfer stimulated by ICOSL, but these did not elicit CD40L transfer even when CD40 was also present in the BSLB. TCR engagement independent IS are stimulated in CD8 effector T cells by NKG2D ligands, but similarly don’t stimulate release of cytolytic effects (36, 37). TCR engagement independent IS-like structures may allow T cells to more closely inspect APC for presence of relevant pMHC. We expect that BSLB will be a useful approach to study synaptic transfer in other biological systems.

We have demonstrated that SE incorporate CD40L as long as CD40 is included in the SLB in addition to a TCR agonist and ICAM-1. This has important implications for SE function in T cell help. CD40L is primarily expressed on CD4^+^ T cells, but low-level mRNA and/or protein expression has also been reported on B cells, basophils, eosonophils, NK cells, macrophages, dendritic cells, smooth muscle cells (38, 39), and platelets (40). Extracellular vesicles from platelets possess CD40L and these vesicles have CD40L dependent adjuvant-like function when injected into mice (40). However, conditional knockout of CD40L in CD4 T cells appears to fully account for the classical defects in antibody class switching and T cell help associated with CD40L deficiency (41). Instead, platelet CD40L appears to function as an integrin ligand in blood clotting such that its physiological function is not CD40 dependent, but integrin β3 dependent (42). In contrast, T cell SEs are generated in an antigen dependent manner and TCR and CD40L are often present in the same SE, thus adding another level of specificity of this post-T cell product. It doesn’t escape our attention that SE may provide an explanation for reports of antigen specific helper factors (43).

T cells transfer CD40L to B cells in an agonist pMHC and CD40 dependent manner (17). We and other investigators speculated that CD40L transfer in extracellular vesicles might be an important component of T cell help (17, 44). Our new results support this speculation in that CD40L is readily incorporated into SE and dSTORM confirms that CD40L and TCR are incorporated in the same CD81 positive vesicles. Gardell and Parker were also able to detect bystander capture of CD40L by CD40 positive, agonist pMHC negative B cells in the presence of CD40 negative, agonist pMHC positive B cells (17). We were unable to recapitulate “bystander” transfer of CD40L when TCR agonists and CD40 are presented on different BSLB. This suggests that active processes in the B cells are required to use CD40 to capture CD40L from a T cell without a TCR agonist to form an IS. A recent study suggests that DC also receive CD40L-CD40 signals without cognate pMHC (45).

Both we (16) and Gardell and Parker (17) utilized anti-CD40L mAb to detect CD40L transfer, which comes with caveats. Since these mAb are function blocking, we naturally were concerned that CD40 in the SLB might compete with the fluorescently tagged anti-CD40L mAb leading to underestimation of transfer. High densities of CD40 in the SLB did suppress the signal with anti-CD40L, but CD40L was still equally depleted from T cells on average, so our measurements of CD40L are likely an underestimate, but over the entire physiological range of CD40 densities we could detect transfer with anti-CD40L mAb. The dSTORM analysis of anti-CD40L localization in SE revealed that CD40L was detected in microclusters distinct from the TCR/ICOS/BST2 microcluster that must correspond to the site of attachment between the SE and the PSLB. This physical segregation of CD40L from the attachment site to the antigen-presenting surface may in part explain its availability to both the anti-CD40L antibody used for detection and CD40 on DCs, which responded functionally to the CD40L on SE. We also noted a CD40/CD40L independent, DC stimulatory activity associated with released SE, which may be explained by recently reported release of mitochondrial DNA associated with T cell exosomes (46), released in parallel with SE.

MS analysis provided a number of new candidates for SE generation and function. We detected many new ESCRT and vesicle trafficking components in SE. We identified two ubiquitin recognizing ESCRT-0 components HRS (Hepatocyte growth factor-regulated tyrosine kinsase substrate; an ESCRT-0 component) and EPN1(47), which may explain why we previously found that HRS is not essential for cSMAC formation (6). We previously could find no role for classical ESCRT-II proteins in SE formation (6), but our MS analysis found ALIX, which bridges TSG101 to ESCRT-III components through its Bro-1 domain (48). Of the candidates investigated by TIRFM, EPN1 was highly concentrated in a ring of discrete punctate structures surrounding the synaptic cleft, which is a pattern observed for other ESCRT components (10). EPN1 may serve in a sorting complex with AP2, clathrin, EPS15 and dynamin (49, 50), which were all detected in SE. We do not know if SE that are transferred through the IS can be released from the recipient APC to engage other APC in a TCR dependent or independent manner. But it is clear that SE attached to CD40 bearing SLB does maintain CD40L that is not engaged by CD40. The MS analysis identified ADAM10 in SE. ADAM10 converts CD40 engaged CD40L into soluble trimers (18), which might release the SE bearing non-CD40 engaged clusters of CD40L to allow interaction with other APC. Further mining of the rich list of proteins from the MS analysis may identify means to selectively interfere with SE formation in vivo to better understand the critical physiological role of CD40L and TCR transfer to APC in synaptic ectosomes.

## Materials and Methods

### Ethics

Leukapheresis products (non-clinical and de-identified) from donor blood were used as a source of human T cells and monocytes. The Non-Clinical Issue division of National Health Service approved the use of leukapheresis reduction (LRS) chambers products at the University of Oxford (REC 11/H0711/7). Clone 35 was isolated from a healthy volunteer where written informed consent was given. Ethical approval was obtained from the University of Oxford Tropical Ethics Committee (OXTREC).

### T cell lymphoblast and Clone 35 culture

CD4^+^ T cell lymphoblasts were generated from human peripheral blood CD4^+^ T cells isolated from healthy donors (51). Briefly, CD4^+^ T cells were isolated by negative selection (RosetteSep Human CD4+ T cell Enrichment Kit, Stemcell technologies) following the manufacturer’s procedure. The CD4^+^ T cells were activated for 3 days using anti-CD3/anti-CD28 T-cell activation and expansion beads (Dynabeads, ThermoFisher Scientific) in complete medium (RPMI 1640 media supplemented with 10% heat-inactivated fetal bovine serum, 50 U/ml of Penicillin-Streptomycin, 2 mM L-Glutamine, 10 mM HEPES, 1 mM Sodium Pyruvate, and 100 μm non-essential amino acids) with 100 U/ml of recombinant human IL-2 (PeproTech), which was replaced every 2 days keeping the cells at a concentration of 1.5 × 10^6^ cells/ml. IL-2 containing media (25 U/mL) was replenished the night before experiments. We refer to these as “T cells” and they were used on day 10 when all division had ceased. The HLA-DRB1*09:01-restricted T cell clone 35 (specific against the influenza H3 HA_338-355_ peptide NVPEKQTRGIFGAIAGFI) were expanded using at a ratio of 1 clone: 2 feeder cells (irradiated, pooled PBMCs from 2-3 healthy donors) at a total cell concentration of 3 × 10^6^ cells/ml in RPMI 1640 supplemented with 10% heat-inactivated AB human serum and 30 μg/ml of PHA for three days. Then, 100 U/ml of recombinant human IL-2 were added to fresh media, which was replaced every 2 days.

### MoDC culture and activation

Monocyte-derived dendritic cells (moDC) were generated from the peripheral blood of healthy adults by first isolating monocytes by negative selection (RosetteSep™ Human Monocyte Enrichment Cocktail, STEMCELL™ technologies) following the manufacturer’s procedure. Then, 1.5-2 × 10^6^ monocytes/ml/cm^2^ were stimulated in complete media supplemented with 100 ng/ml of recombinant human GM-CSF and 200 ng/ml of recombinant human IL-4 (52). After 5 to 7 days of culture, moDC were used in experiments.

### Antibodies

Primary monoclonal antibodies (mAb) used for dSTORM were anti-TCR-Alexa Fluor^®^ (AF) 488 (clone IP26; BioLegend), anti-CD40L-AF647 (clone 24-31; BioLegend), anti-ICOS-AF647 (clone C398.4A; BioLegend), anti-BST2-AF647 (clone RS38E; BioLegend), anti-HLA-DR-AF488 (clone L243; BioLegend), anti-CD81-AF647 (clone 5A6; BioLegend) anti-CD83-AF647 (clone HB15e; BioLegend) and Wheat Germ Agglutinin WGA-CF568 (Biotum) to label the surface of the SEs. All antibody clones used to assess relative or absolute quantification of protein transfer from cells to BSLB are listed in Table S1. Isotype controls matching the relevant fluorescent dyes were used for background correction and gating. Other mAb or affinity purified antibodies are described with specific methods below.

### Small unilamellar vesicles (SUVs)

SUV are defined as vesicles in the 20-100 nm range. SUV were formed by extrusion as described using the Avanti Miniextruder with a 100 nm filter (53). When SUV were used to mimic SE, all lipids were combined prior to SUV formation, whereas BSLB and PSLB composition could be determined by mixing different proportions of stock SUVs as the final bilayer composition is determined by the average of the input SUV. NTA-SUVs for attachment of His tagged proteins were composed of 85.5 mol% DOPC, 2 mol% head group labeled ATTO-390-DOPE, and 12.5 mol% DOGS-NTA at a total lipid concentration of 4 mM. Plain SUVs that were not able to bind His tagged proteins, were composed of 98 mol% DOPC and 2% ATTO-390-DOPE at a total lipid concentration of 4 mM. Stock SUV for formation of BSLB or PSLB were composed of 0.4 mM solution of lipids in PBS with 100 mol% DOPC; 75 mol% DOPC and 25 mol% DOGS-NTA; 98 mol% DOPC and 2 mol% DOPE-CAP-Biotin; or 98% DOPC; 2 mol% ATTO-(390 or 488)-DOPE. These stocks could be mixed in different ratios prior to formation of BSLB or PSLB to generate mobile bilayers of the desired final composition. All lipids were purchased from Avanti Polar Lipids, Inc (Alabaster, AL).

### Nanoparticle Tracking Analysis

A 10 μL aliquot of SUVs or eluted SE preparation was re-suspended in PBS in a 1:100 dilution and kept on ice for Nanoparticle Tracking Analysis (NTA). The instrument used for NTA was Nanosight NS300 (Malvern Instruments Ltd) set on light scattering mode and instrument sensitivity of 15. Measurements were taken with the aid of a syringe pump to improve reproducibility. Three sequential recordings of 60 seconds each were obtained per sample and NTA 3.2 software was used to process and average the three recordings to determine the mean size.

### moDC activation by CD40L on SUV

His-tagged recombinant soluble CD40L (sCD40L, BioLegend) was incubated with NTA-SUV or plain SUV, at ratios designed to match CD40L densities found on SE for 20 min at 24°C prior to addition to the moDCs. After 24 h, moDCs were recovered by spinning down plates at 1500 rpm for 5 min and resuspended in flow cytometry staining buffer (10% Heat-Inactivated Goat Serum, 0.04% sodium azide in PBS pH 7.4) and incubated for 30 min at 4ºC. A final concentration of 10-30 nM of each mAb was used. The multicolor panel included anti-HLA-DR PerCP (clone L243), anti-CD40 AF647 (clone 5C3), anti-ICAM-1 Brilliant Violet 510/Brilliant Violet 785 (clone HA58), anti-CD80 PE (clone 2D10), anti-CD86 Brilliant Violet 785 (clone IT2.2) and anti-ICOSL PE-Cy7 (clone 2D3). Isotype control antibodies clones MOPC-21 (IgG1, κ), MOPC-173 (IgG2a, κ) and MPC-11 (IgG2b, κ) were used matching the relevant fluorescent dyes. Staining was performed for 30 min at 4 ºC in the dark and constant agitation after which cells were washed twice and single cell fluorescence measurements were made by flow cytometry.

### Bead Supported Lipid Bilayers

Silica beads (5.0 μm diameter, Bangs Laboratories, Inc.) were washed extensively with PBS in a 1.5 ml conical microcentrifuge tubes. BSLBs were formed by incubation with mixtures of SUVs to generate a final lipid composition of 0.2 mol% ATTO 488-DOPE; 12.5 mol% DOGS-NTA and a mol% of DOPE-CAP-Biotin to yield 10-5000 molecules/μm^2^ UCHT1-Fab in DOPC at a total lipid concentration of 0.4 mM. The resultant BSLB were washed with 1% human serum albumin (HSA)-supplemented HEPES-buffered saline (HBS), subsequently referred to as HBS/HSA. After blocking with 5% casein in PBS containing 100 μM NiSO_4_, to saturate NTA sites, 50 μg/mL unlabelled streptavidin was then coupled to biotin head groups by incubation with concentrations of streptavidin determined to yield 10-5,000 molec. /μm^2^ site densities. After 20 minutes, the BSLB were washed 2× with HBS-HSA and biotinylated UCHT1-Fab (variable density as indicated), His-tagged ICAM-1 (200 molec. /μm^2^), CD40 (500 molec./μm^2^), and ICOSL (100 molec./μm^2^) were then incubated with the bilayers at concentrations to achieve the indicated site densities (in range of 1-100 nM). Excess proteins were removed by washing with HBS/HSA after 20 minutes. T cells (5×10^5^/well) were incubated with BSLB at 1:1 ratio in a V-bottomed 96 well plate (Corning) for 1 hr at 37ºC in 100 μl HBS/HSA. BSLB: cell conjugates were pelleted at 500 × g for 1 minute prior to resuspension in 50 mM EDTA in PBS at 4ºC to release His-tagged proteins from the BSLB, while leaving the UCHT1-Fab attached, thus selectively retaining TCR^+^ SE. The single BSLB and cells were gently resuspended prior to staining for flow cytometry analysis or sorting.

### Calibration of flow cytometry data

T cells and BSLB were analyzed using antibodies with known AF647:Ab ratio (Table S1) in parallel with the Quantum AF647 Molecules of Equivalent Soluble Fluorescent dye (MESF) beads, allowing the calculation of the absolute number of mAb bound per T cell and per BSLB after subtraction of unspecific signals given by isotype control antibodies.

### Airyscan microscopy

Airyscan imaging of BSLB-cell conjugates was performed on a confocal laser-scanning microscope Zeiss LSM 880 equipped with Airyscan detection module (Zeiss, Oberkochen, Germany) using the Plan-Apochromat 63×/1.46 Oil objective (Zeiss, Oberkochen, Germany). The Argon laser at 488 nm and diode laser at 561 nm were used as excitation sources, with power setting of ~1% and ~6%, respectively, which is equivalent to 1 mW and 10 mW. The powers were set in this range in order to achieve the comparable strength of fluorescent signal for both channels. Fluorescence emission was collected at around 515 nm and 653 nm for the green and magenta channels, respectively, with the following filters BP420-480+BP495-550 (green) and BP555-620+LP645 (magenta). The emission signals were collected on the 32 channel GaAsP-PMT Airy detector. The datasets were acquired as Z-stacks with 43.5 nm pixel size and 185 nm axial steps, which correspond to ~50-55 slices per 3D data set. ZEN Airyscan software (Zeiss) was used to process the acquired data sets. This software processes each of the 32 Airy detector channels separately by performing filtering, deconvolution and pixel reassignment in order to obtain images with enhanced resolution and improved signal to noise ratio. The value of Wiener filter in ZEN software was chosen in accordance with the value in “auto” reconstruction modality and was set around 7, to ensure the absence of deconvolution artefacts (54). Drift was corrected using the MultiStackReg plug-in of ImageJ (National Institute of Health). Rendering was performed in Imaris software (Bitplane).

### Planar Supported Lipid Bilayers (PSLB)

SUV mixtures were injected into flow chambers formed by sealing acid piranha cleaned glass coverslips to adhesive backed plastic manifolds with 6 flow channels (StickySlide VI 0.4; Ibidi) (16). After 20 minutes the channels were flushed with HBS-HSA without introducing air bubbles to remove excess SUVs. After blocking for 20 min with 5% casein supplemented with 100 μM NiCl_2_, to saturate NTA sites, followed by 15 min incubation with streptavidin (Sigma Aldrich), washing and then monobiotinyated or His-tagged proteins were incubated on bilayers for additional 20 min. Protein concentrations required to achieve desired densities on bilayers were calculated from calibration curves constructed from flow-cytometric measurements of BSLB, compared with reference beads containing known numbers of the appropriate fluorescent dyes (Bangs Laboratories). Bilayers were continuous liquid disordered phase as determined by fluorescence recovery after photobleaching with a 10 μm bleach spot on an FV1200 confocal microscope (Olympus).

### T cell Immunological Synapse formation on PSLB

CD4^+^ T cells were incubated at 37°C on SLB containing either ICAM-1 alone, ICAM-1 and UCHT1-Fab or ICAM-1 UCHT1-Fab, CD40 and ICOSL. After 20-90 minutes of incubation the cells either fixed with 4% electron microscopy grade formaldehyde in PHEM buffer (10 mM EGTA, 2 mM MgCl_2_, 60 mM Pipes, 25 mM HEPES, pH 7.0), permeabilized with 0.1% Triton X-100 (if necessary for access to intracellular spaces) and stained with primary conjugated antibodies and imaged. Alternatively, the cells were washed off with cold PBS and the SE left behind were stained with directly conjugated antibodies and fixed with 4% formaldehyde in PHEM buffer. Prior the labelling of moDCs with mAbs on SLB for TIRF imaging, the cells were blocked for Fc receptors with 5% HSA and 5% goat or donkey serum for 1 h at 24°C.

### Total internal reflection fluorescence microscopy (TIRFM)

TIRFM was performed on an Olympus IX83 inverted microscope equipped with a 4-line (405 nm, 488 nm, 561 nm, and 640 nm laser) illumination system. The system was fitted with an Olympus UApON 150 × 1.45 numerical aperture objective, and a Photomertrics Evolve delta EMCCD camera to provide Nyquist sampling. Live experiments were performed with an incubator box maintaining 37°C and a continuous autofocus mechanism. Quantification of fluorescence intensity was performed with ImageJ (National Institute of Health).

### dSTORM imaging and data analysis

For three colour dSTORM imaging extracellular vesicles were stained using either wheat germ agglutinin (WGA) directly conjugated with CF568 (Biotium) or anti-CD81-AF647. First, 640-nm laser light was used for exciting the AF647 dye and switching it to the dark state. Second, 488-nm laser light was used for exciting the AF488 dye and switching it to the dark state. Third, 560-nm laser light was used for exciting the CF568 dye and switching it to the dark state. An additional 405-nm laser light was used for reactivating the AF647, AF488 and CF568 fluorescence. The emitted light from all dyes was collected by the same objective and imaged onto the electron-multiplying charge-coupled device camera with an effective exposure time of 10 ms. A maximum of 5,000 frames for antibodies conjugated with AF647, CF568 and AF488 condition were acquired. For visualizing the WGA labelled extracellular vesicles minimum of 80,000 frames were acquired. For each receptor, the specificity of the labelling was confirmed by staining the vesicles with isotype-matched control antibodies (data not shown).

Because multicolour dSTORM imaging is performed in sequential mode by using three different optical detection paths (same dichroic but different emission filters), an image registration is required to generate the final three-color dSTORM image (55–57). Therefore, fiducial markers (TetraSpek Fluorescent Microspheres; Invitrogen) of 100 nm, which were visible in 488-nm, 561-nm and 640-nm channels, were used to align the 488-nm channel to 640-nm channel. The difference between 561-nm channel and 640-nm channel was negligible and therefore transformation was not performed for 561-nm channel. The images of the beads in both channels were used to calculate a polynomial transformation function that maps the 488-nm channel onto the 640-nm channel, using the MultiStackReg plug-in of ImageJ (National Institute of Health) to account for differences in magnification and rotation, for example. The transformation was applied to each frame of the 488-nm channel. dSTORM images were analyzed and rendered as previously described (58, 59) using custom-written software (Insight3, provided by B. Huang, University of California, San Francisco). In brief, peaks in single-molecule images were identified based on a threshold and fit to a simple Gaussian to determine the x and y positions. Only localizations with photon count ≥ 2000 photons were included, and localizations that appeared within one pixel in five consecutive frames were merged together and fitted as one localization. The final images were rendered by representing the x and y positions of the localizations as a Gaussian with a width that corresponds to the determined localization precision. Sample drift during acquisition was calculated and subtracted by reconstructing dSTORM images from subsets of frames (500 frames) and correlating these images to a reference frame (the initial time segment). ImageJ was used to merge rendered high-resolution images (National Institute of Health).

### CBC analysis

Coordinate-based colocalization (CBC) mediated analysis between two proteins was performed using an ImageJ (National Institute of Health) plug-in (60) based on an algorithm described previously (27). To assess the correlation function for each localization, the x-y coordinate list from 488-nm and 640-nm dSTORM channels was used. For each localization from the 488-nm channel, the correlation function to each localization from the 640-nm channel was calculated. This parameter can vary from −1 (perfectly segregated) to 0 (uncorrelated distributions) to +1 (perfectly colocalized). The correlation coefficients were plotted as a histogram of occurrences with a 0.1 binning. The Nearest-neighbor distance (NND) between each localization from the 488-nm channel and its closest localization from the 640-nm channel was measured and plotted as the median NND between localizations per cell.

### Cross-correlation analysis

Cross correlation analysis is independent of the number of localizations and is not susceptible to over-counting artifacts related to fluorescent dye re-blinking and the complements other approaches (28). Cross-correlation analysis between two proteins was performed using MATLAB software provided by Sarah Shelby and Sarah Veatch from University of Michigan. Regions containing cells were masked by region of interest and the cross-correlation function from x-y coordinate list from 488-nm and 640-nm dSTORM channels was computed from these regions using an algorithm described previously (28, 61, 62). Cross-correlation functions, C(r,q), were firstly tabulated by computing the distances between pairs of localized molecules, then C(r) is obtained by averaging over angles. Generally, C(r) is tabulated from ungrouped images, meaning that localizations detected within a small radius in sequential frames are counted independently. Finally, a normalized histogram with these distances was constructed into discrete bins covering radial distances up to 1000 nm. Cross-correlation functions only indicate significant correlations when the spatial distribution of the first probe influences the spatial distribution of the second probe, even when one or both of the probes are clustered themselves. Error bars are estimated using the variance within the radial average of the two dimensional C(r, q), the average lateral resolution of the measurement, and the numbers of probes imaged in each channel. The cross-correlation function tabulated from the images indicates that molecules are highly colocalized, where the magnitude of the cross-correlation yield (C(r) > 1) is higher than randomly co-distributed molecules (C(r) = 1).

### Cytokine array

Primary moDCs were incubated on the extracellular vesicles derived from CD4^+^ T cells on SLB containing either only ICAM-1 (200 molec/μm^2^), ICAM-1 and UCHT1-Fab (300 molec/μm^2^), or ICAM-1, UCHT1-Fab, CD40 (500 molec/μm^2^) and ICOSL (100 molec/μm^2^), at 37°C for 24 h. Cell supernatants were recovered and centrifuged at 350 g for 5 min at RT to remove cells and cell debris. Cytokine production was quantified in the supernatants by Human XL Cytokine Array kit (ARY022B; R&D Systems), according to manufacturer’s instructions. The positive signal from cytokines was determined by measuring the average signal of the pair of duplicate spots by using ImageJ (National Institute of Health). Differences between arrays were corrected by using the average intensity of positive spots within the array. Fold change of the cytokine production between conditions was determined by normalizing the data to SLB containing only ICAM-1.

### Mass Spectrometry

AF488+ BSLB were sorted on a FACS ARIA III and lysed by sonication (Bioruptor Pico) in 0.5% NP-40 in 50 mM ammonium bicarbonate and 6 M urea. Cysteines were reduced and alkylated by addition of first 5 μl of 200 mM dithiothreitol (30 minutes at 24°C) and 10 μl of 200 mM iodoacetamide (60 minutes at RT in dark). The protein solution was then precipitated with chloroform and methanol (63), and resuspended in 6 M Urea. For digest the protein solution was diluted in 50 mM ammonium bicarbonate, pH 7, and 0.6 μg trypsin was added for digest at 37°C overnight. Peptides were desalted with a C18 solid phase extraction cartridge (SOLA, Thermo Fisher Scientific) and resuspended in 15 μl 2% acetonitrile and 0.1% trifluoroacetic acid in water. Samples were analyzed on a LC-MS/MS platform consisting of Orbitrap Fusion Lumos coupled to a UPLC ultimate 3000 RSLCnano (both Thermo Fisher Scientific). Samples were loaded in 1% acetonitrile and 0.1% trifluoroacetic acid in water and eluted with a gradient from 2 to 35 % acetonitrile, 0.1% formic acid and 5% dimethylsulfoxide in water in 60 minutes with a flow rate of 250 nl/min on an EASY-Spray column (ES803, Thermo Fisher Scientific). The survey scan was acquired at a resolution of 120.000 between 380-1500 m/z and an automatic gain control target of 4E5. Selected precursor ions were isolated in the quadrupole with a mass isolation window of 1.6 Th and analyzed after CID fragmentation at 35% normalized collision energy in the linear ion trap in rapid scan mode. The duty cycle was fixed at 3 seconds with a maximum injection time of 300 ms, AGC target of 4000 and parallelization enabled. Selected precursor masses were excluded for the following 60 seconds. Proteomic data was analyzed in Maxquant (V1.5.7.4, ref) using default parameters and Label Free Quantitation. The data was searched against the mouse canonical Uniprot database (29/07/2015) and the human Uniprot database (15/10/2014). FDR on peptide and protein level were set to 1%. Second peptide and “match between runs” options were enabled.

The MS data set is available to reviewers prior to publication:

ProteomeXchange Consortium via the PRIDE (64) partner repository with the dataset identifier PXD007988

Upon publication the dataset will be publically available.

### Statistical analysis

All statistical analyses were performed using SigmaPlot 13.0 (Systat Software Inc), OriginPro 2017 software (OriginLab) or GraphPad Prism v 7.0 and 8.0 (GraphPad Software, Inc.). Statistical analyses are detailed in each figure legend.

## Acknowledgments

We thank E. Kurz, A. Afrose, H. Rada and P. Hernandez-Varas for technical assistance; A. M. Santos, S. J. Davis, S. Campion, P. Borrow and P. Lipsky for helpful suggestions; and A. Kvalvaag, P. Borrow, C. Vinuesa and G. Victora for critical reading. Supported by the ERC AdG 670930 (MLD), the Wellcome Trust PRF 100262 (MLD), the Kennedy Trust (to MLD and BMK), the NIH AI043542 (MLD), the NIH tetramer core facility, the EMBO ALTF 1420-2015 (PFCD), Medical Research Council (YCD and TD), UK and Chinese Academy of Medical Sciences Innovation Fund 2018-I2M-2-002 (YCD and TD) and Cancer Research UK A19277 (EO).

## Author Contributions

MLD, DGS, PFCD, SB and EBC conceived studies, performed experiments, interpreted data and wrote paper. SV and VM provided key infrastructure. KK acquired and rendered Airyscan images. YP and TD provided and validated clone 35 cells. MT and EO provided Nanosight data and interpretation. SB, RF and BMK-acquired, analyzed and interpreted Mass Spectroscopy data.

## Supplementary Materials

**Figure 2 – Figure supplement 1.**
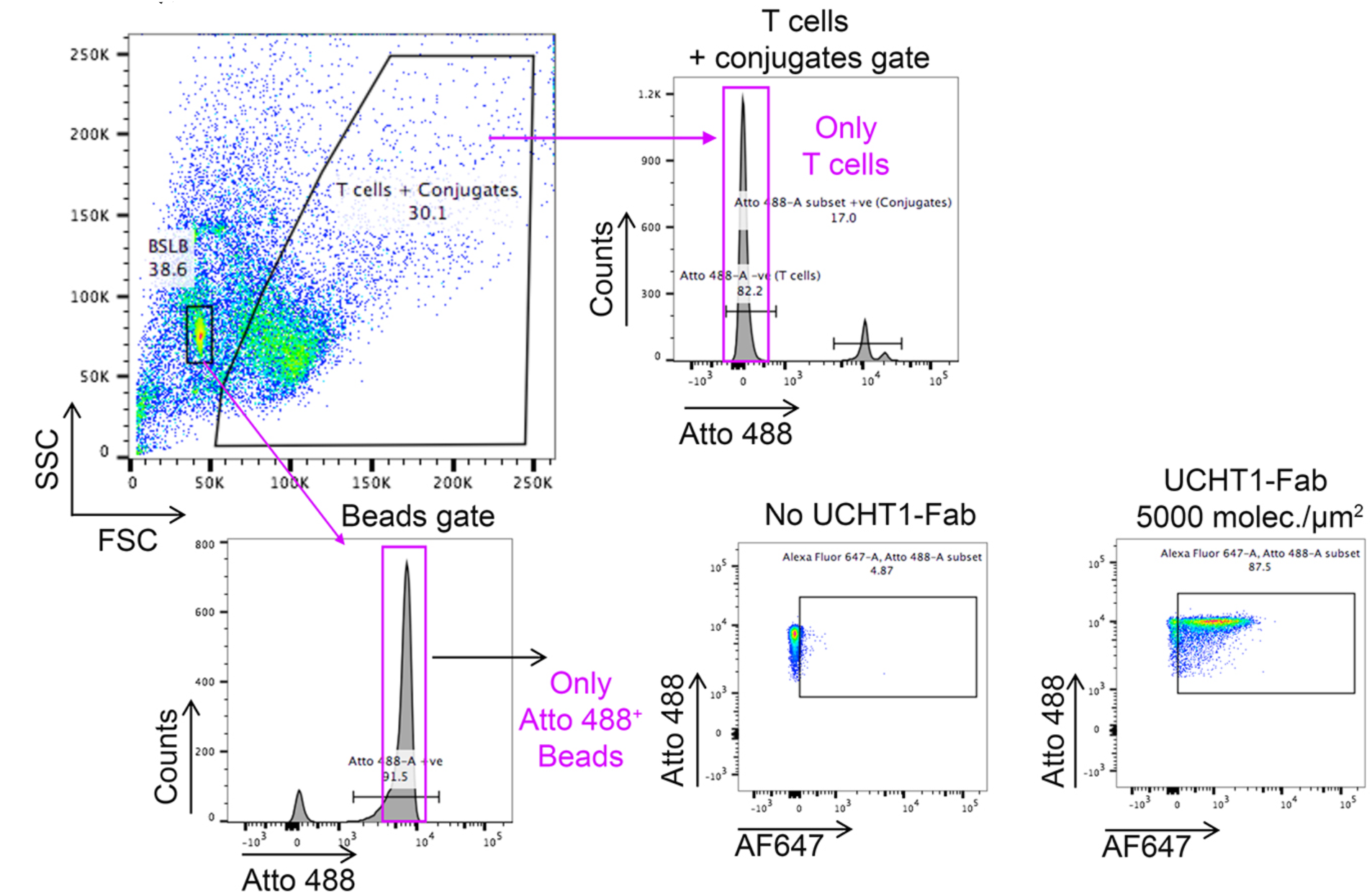
Gating Strategy for SE capture on BSLB. Forward/Side site scatter plot of beads and T cells following liberation of BSLB from with EDTA/PBS. Beads have a distinctive and narrow scatter signature, but this partly overlaps with cell debris. Beads are thus also positively discriminated from cell debris using Atto-488 lipids in the BSLB. Cells and conjugates are discriminated based on Atto-488-ve signal for cells and a combination of Atto-488+ve signal and higher light scattering signals for conjugates.

**Figure 2 – Figure supplement 2.**
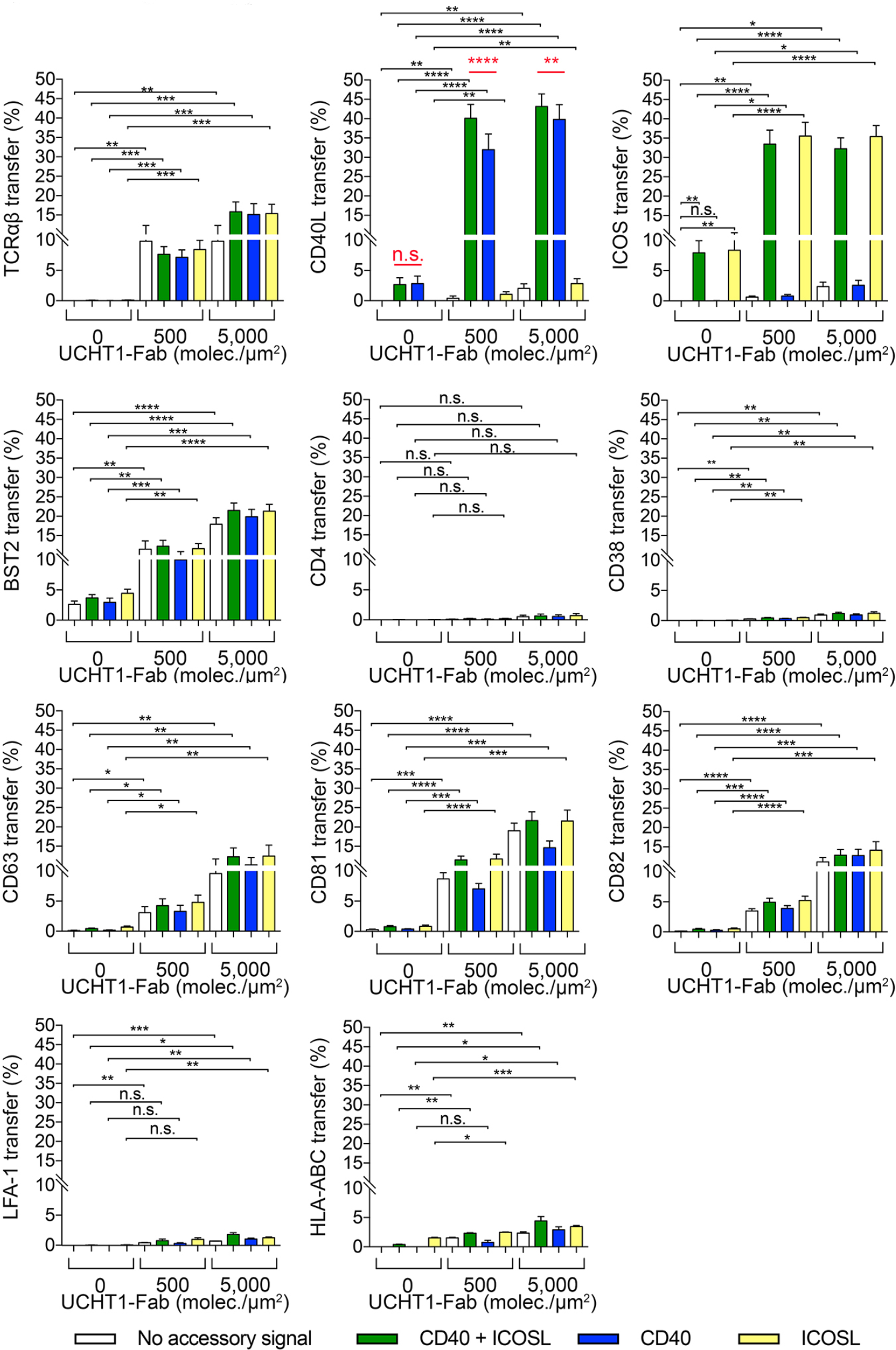
ICOS significantly increases CD40L transfer to BSLB. Per marker statistical analyses of heat maps shown in Figure 2D. *P* values <0.05 were considering significant. Repeat Measures ANOVA with Geisser-Greenhouse correction was performed. Significance was calculated comparing in each bilayer composition group (either No accessory signal, CD40 + ICOSL, CD40 or ICOSL) at 500 and 5,000 molec./μm^2^ of UCHT1-Fab to No UCHT1-Fab. *P* values: *<0.05, **<0.005; ***<0.0005; ****<0.0001; n.s.= non-significant.

**Figure 2 – Figure supplement 3.**
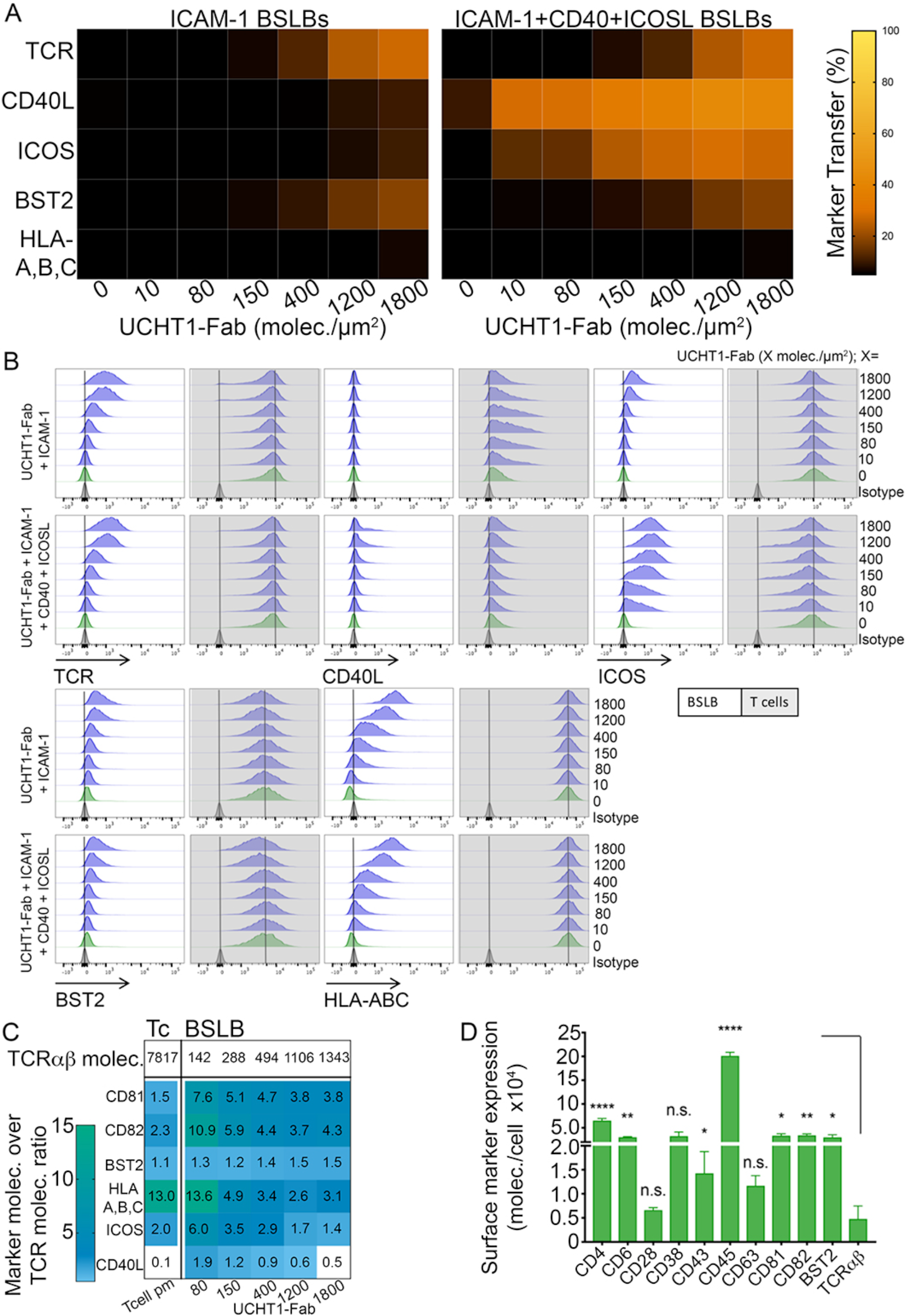
Efficient transfer of CD40L to BSLB at low UCHT1-Fab densities. (A) Percent of protein transfer (%) from human T cells to BSLBs containing increasing densities of UCHT1-Fab (0-1800 molec./μm^2^) and two different BSLB compositions: UCHT1-Fab + ICAM-1 (left) or UCHT1-Fab +ICAM-1 + CD40 + ICOSL (right). (B) Representative histograms showing as in A the synaptic transfer of proteins from T cells (grey background half offset histograms) to BSLBs (white background half-offset histograms) across increasing densities of UCHT1-Fab. (C) Molecules transferred to BSLB per molecule of TCRαβ transferred (marker to TCRαβ ratio). The first row depicts the mean absolute number of TCRαβ molecules per cell (first column) or per BSLB according to different UCHT1-Fab densities (columns 2-6) as reference of the number of molecules used in the calculation of molecular ratios (rows 2-7). Except for BST2, all the other markers reduce their transfer as the transfer of TCRαβ increases. The increased ratios of markers on BSLBs compared to the T cell PM is another form to depict the degree of protein enrichment in SE, but compared to the normal protein to TCRαβ ratio found on the PM of a non-activated T cell (shown are ratios for the mean values from 6 different donors). (D) CD4^+^ T cells were stained for the absolute quantification of surface markers using antibodies with known fluorescent dye to protein (F/P) ratios (See Table S1). For analyses, dead cells were excluded with Propidium Iodide (0.5 μg/mL) during acquisition. Final molecules per cell were obtained by dividing the molecules of equivalent soluble fluorescent dye value (MESF/cell; interpolated from background-corrected GMFI values) by the F/P value of the relevant antibody. *P* values < 0.05 (*); <0.002 (**); <0.0002 (***); <0.0001 (****) were considered significant

**Figure 2 – Figure supplement 4.**
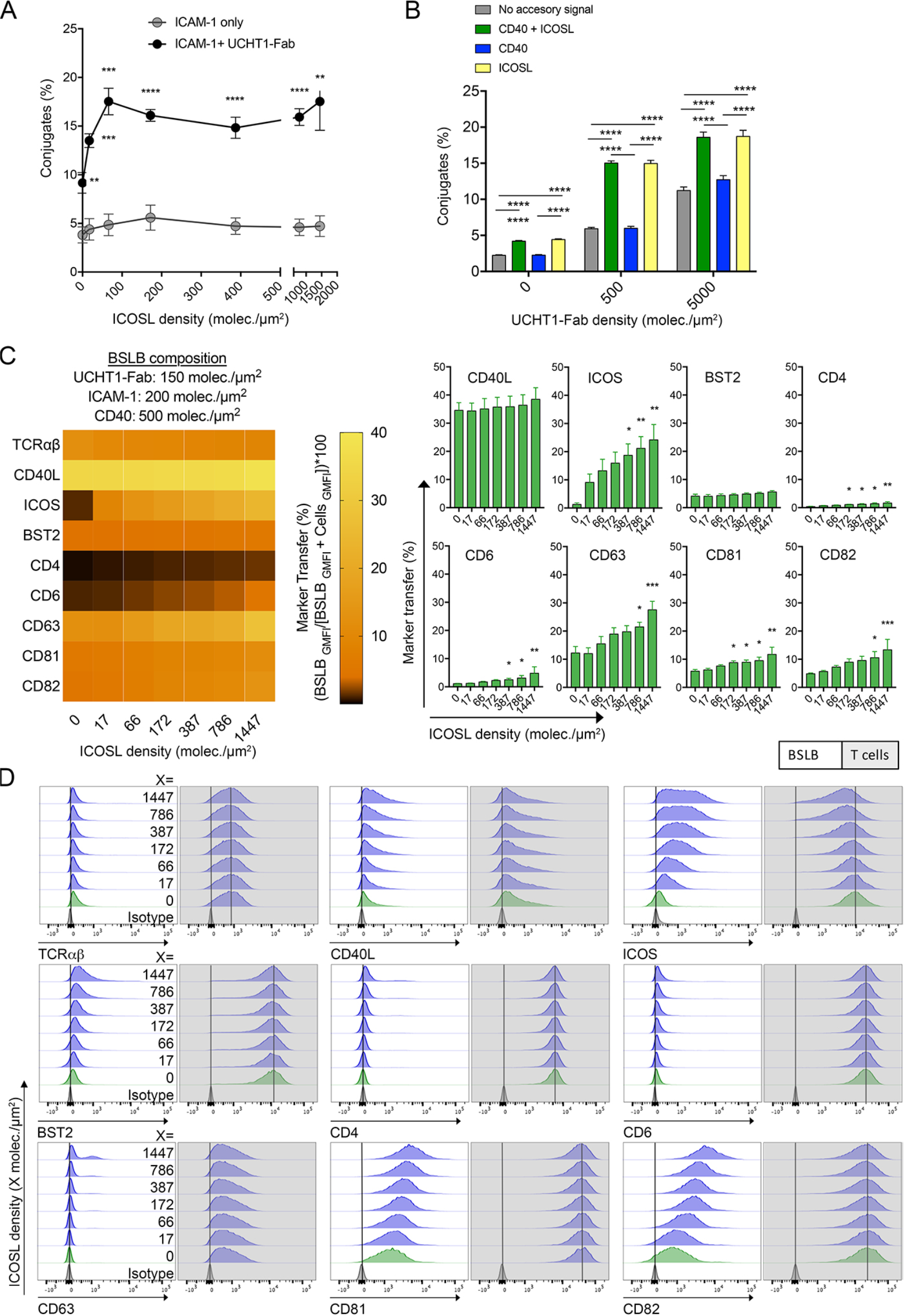
ICOSL increases T cell: BSLB conjugate formation and ICOS transfer. (A) % T cell: BSLB conjugate formation in response to increasing densities of ICOSL. (B) % T cell: BSLB conjugate formation in response to increasing densities of UCHT1-Fab in the presence or absence of accessory proteins ICOSL and or CD40. Multiple t test with false discovery rate of 1%. (C) Using a multicolor flow cytometry panel, the relative transfer of proteins (% of transfer) in response to increasing densities of ICOSL was determined. T cells were stimulated with BSLBs containing 200 molec./μm^2^ of ICAM-1, 150 molec./μm^2^ of UCHT1-Fab and 500 molec./μm^2^ of CD40. After 1h, cell: bead conjugates were dissociated with cold 50 mM EDTA-PBS, and then stained with a multicolor panel using the same antibody clones described in Table S1. We used CD4 as a control protein whose relative (% of total CD4 signal) transfer was not enriched in our single-color experiments (see Figure 2D). Each heat map square represent mean +/-SEM of data collected from 5 donors across 2 independent experiments. Shapiro-Wilk normality test. Non-Normally distributed data was analyzed using no matching or pairing Kruskal-Wallis test with Dunn’s multiple comparisons to the rank of conditions with no ICOSL. Normally distributed data was analyzed using one-way ANOVA with Holm-Sidak’s multiple comparison test to the mean of the no ICOSL group. P < 0.03 was considered significant. *<.03; **<.002; ***<.0002; ****<.0001. (D) Representative off set histograms of BSLBs (white background) and T cells (grey background) analyzed after dissociation of conjugates. Histograms for controls stained with antibodies of appropriate isotype conjugated with the relevant fluorescent dye are shown in grey.

**Figure 2 – Figure supplement 5.**
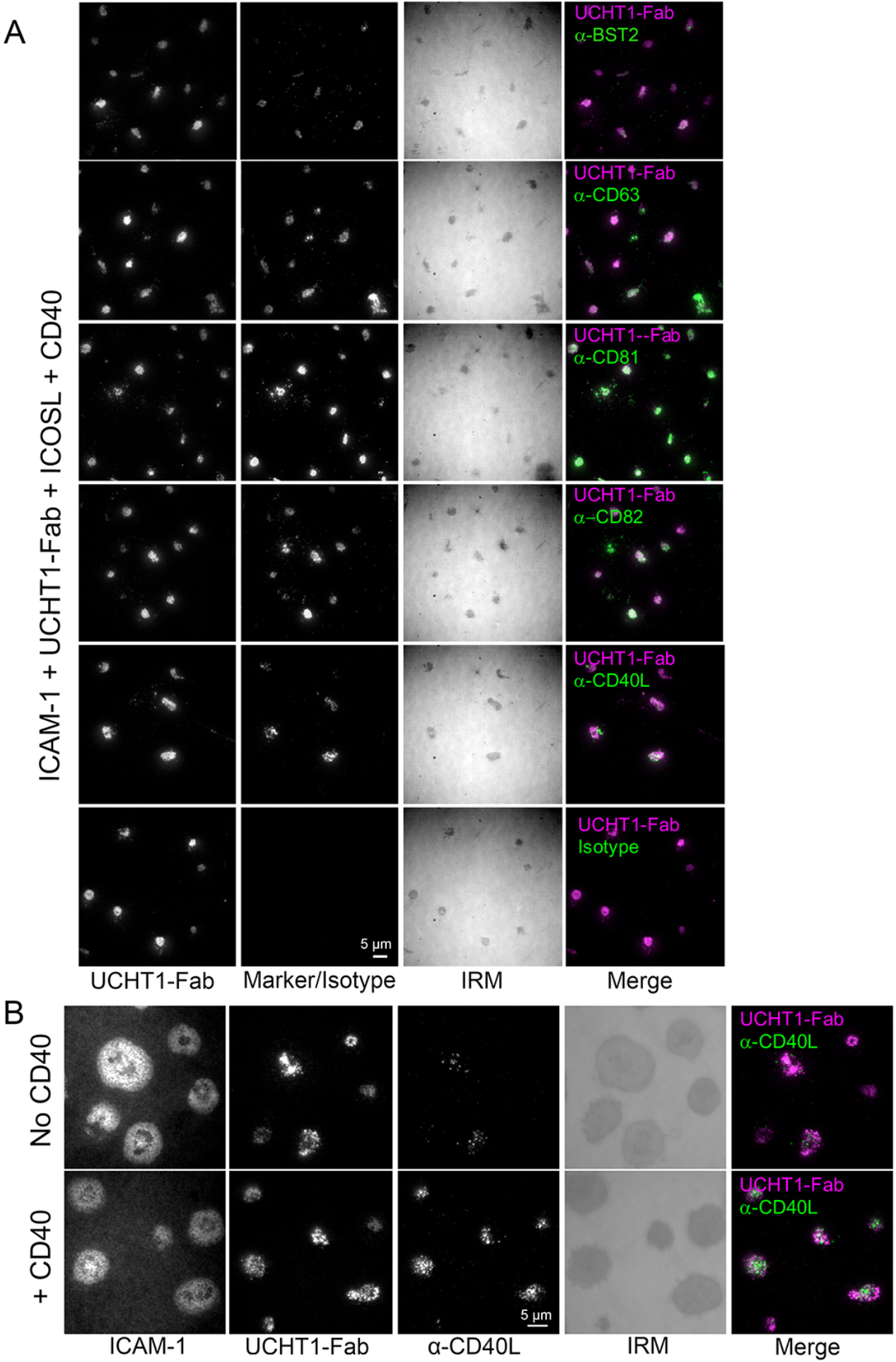
BST2, CD63, CD81 and CD82 and CD40L localize to the synaptic cleft of UCHT-1 Fab stimulated cells. (A) Representative TIRF and IRM images showing that proteins characterized by flow cytometry as synaptically transferred and deposited on PSLB containing ICAM-1, UCHT1-Fab, CD40 and ICOSL. T cells were incubated for 1h at 37ºC and 5% atmospheric CO_2_, then cells were stained for 20 min at RT with anti-protein AF647 antibodies (5 μg/mL each). After two washes, cells were fixed with 4% PFA in PHEM buffer and imaged by TIRFM. (B) Representative TIRF and IRM images showing CD40L stained extracellular vesicles released by CD4^+^ T cells incubated for 15 minutes on PSLB coated with either ICAM-1 with UCHT1-Fab (top panels) or ICAM-1 with UCHT1-Fab and CD40 (bottom panels). Cells were incubated on the PSLB in the presence of anti-CD40L clone 24-31 fluorescently labeled with Alexa Fluor^®^ 647.

**Figure 7 – Figure supplement 1.**
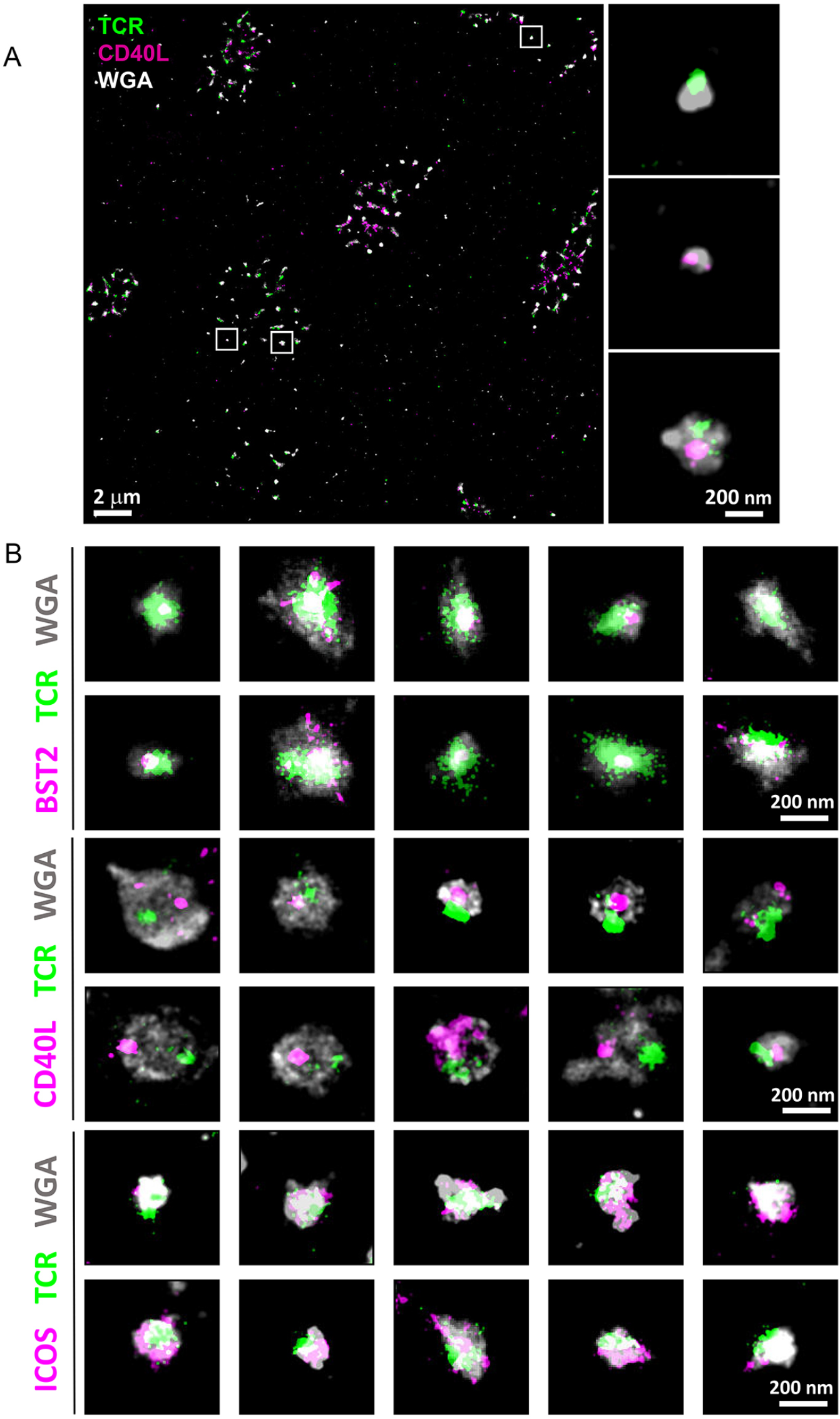
TCR co-localization with BST2 and ICOS and segregation from CD40L. (A) Representative dSTORM images showing TCR (green) and CD40L (magenta) on WGA labeled SE (gray). Insets show examples of SEs containing only TCR, only CD40L or both proteins. (B) Multiple examples of SEs released by CD4^+^ T cells incubated for 90 min on PSLB coated with ICAM-1, UCHT1-Fab, CD40 and ICOSL. The SEs were stained with anti-TCRαβ-AF488, WGA-CF568 to visualize the lipid membrane and with anti-CD40L-AF647, anti-ICOS-AF647 or anti-BST2-AF647.

**Figure 7 – Figure supplement 2.**
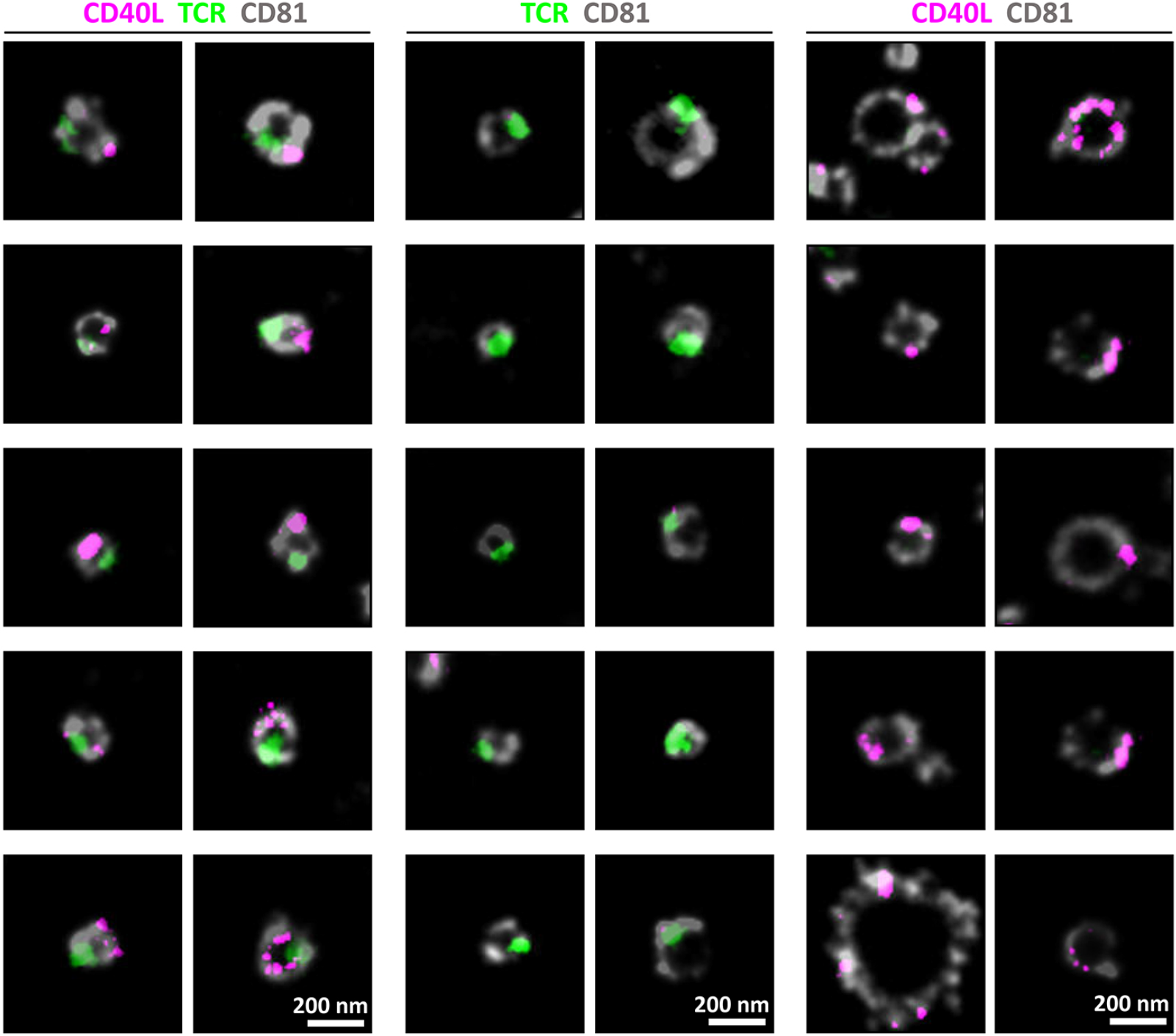
CD81 co-localization with TCR and CD40L. Multiple examples of SEs released by CD4+ T cells incubated for 90 min on PSLB coated with ICAM-1, UCHT1-Fab, CD40 and ICOSL. The SEs were stained with anti-TCRαβ-AF488, anti-CD81-AF568 to visualize the SE membrane and with anti-CD40L-AF647.

**Figure 7 – Figure supplement 3.**
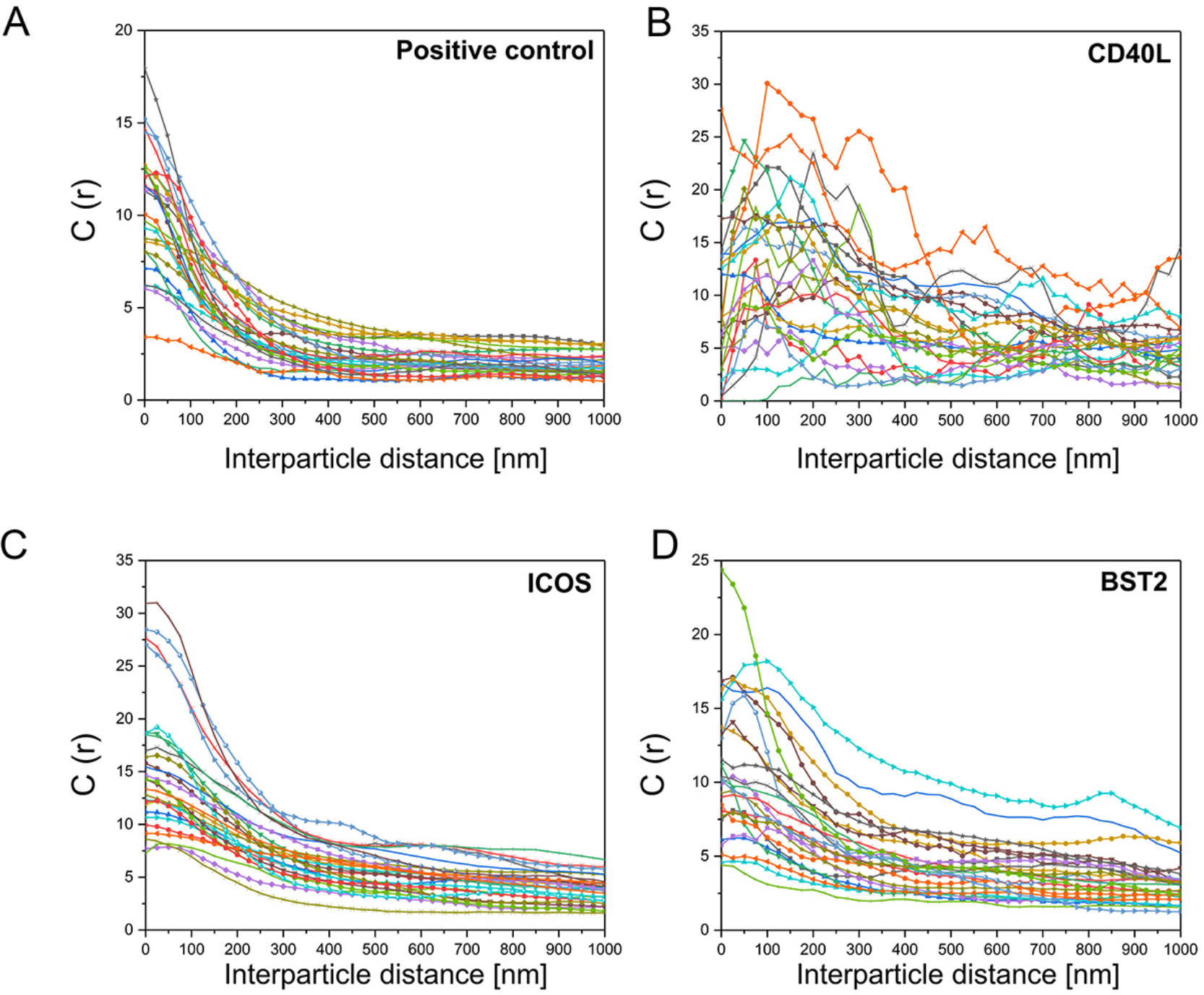
Distinct cross-correlation distances for TCR with BST2 and ICOS versus CD40L. Examples of cross-correlation analysis for positive control data (A), between TCR and CD40L (B), TCR and ICOS (C), or between TCR and BST2 (D) from multiple cells.

**Figure 8 – Figure supplement 1.**
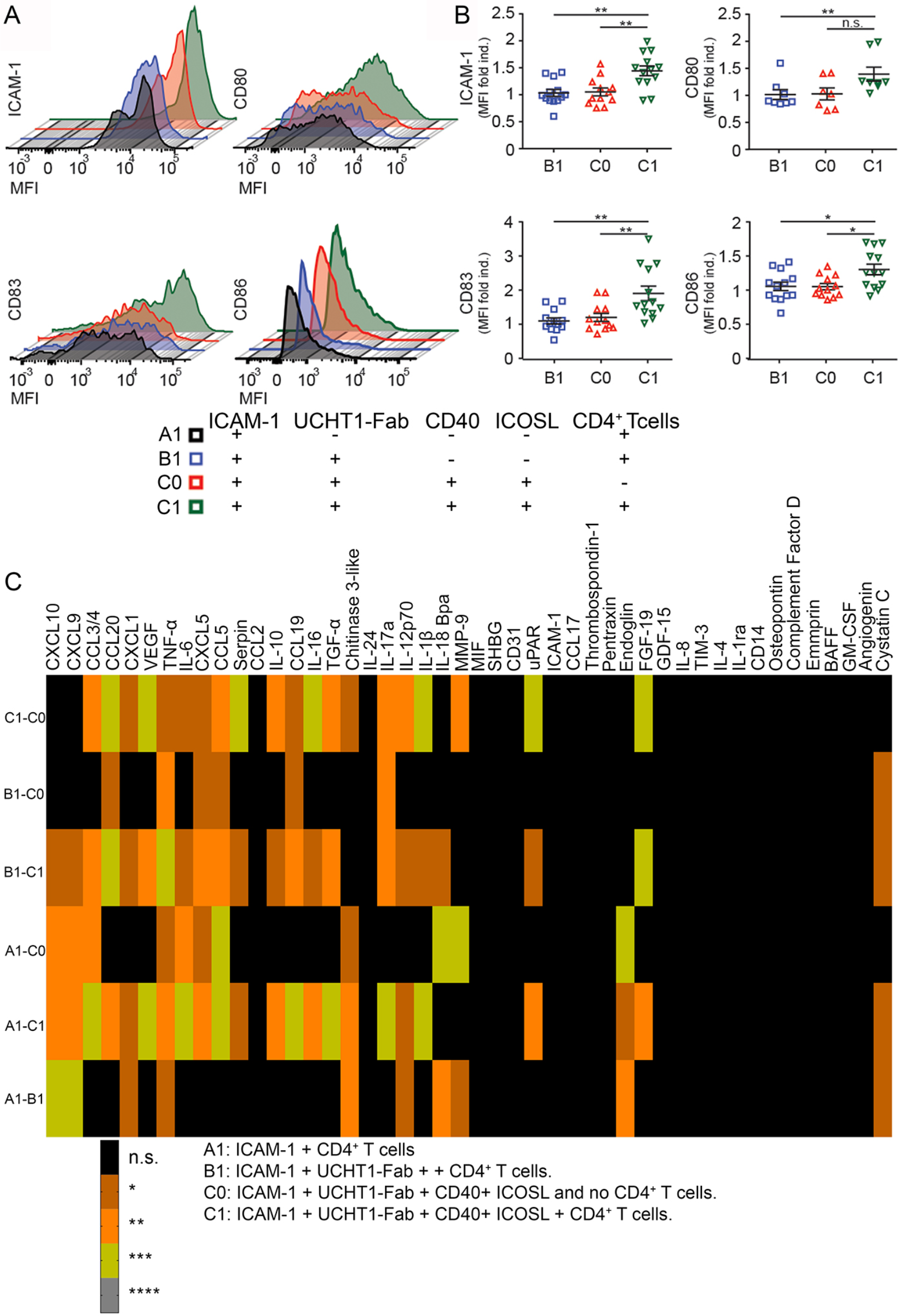
SE captured on PSLB efficiently activate moDCs. (A) Overlaid histograms illustrating the expression of ICAM-1, CD80, CD83 and CD86 maturation markers on DCs following 24h of stimulation on PSLBs loaded with SEs derived from CD4+ T cells. SE-coated PSLB were obtained after the removal of T cells from bilayers containing different stimuli (either A1, B1, C1 or C0). (B) Distribution of ICAM-1, CD80, CD83 and CD86 expression levels in DCs as shown in A. Each symbol represents a donor. (C) Statistical significances for the cytokines/chemokines from Figure 8C. ns, not significant; *, p ≤ 0.05; **, p ≤ 0.01; ***, p ≤ 0.001; ****, p ≤ 0.0001, Kruskal-Wallis with Newman–Keuls post-hoc test.

**Table S1.** Summary of antibodies used for determination of relative and absolute enrichment of bead-transferred proteins.

| Antibody (anti-human) | Clone | Isotype | Mean F/P |
| --- | --- | --- | --- |
| TCR$\alpha\beta$ | IP26 | Mouse IgG1, $\kappa$ | 4.20 |
| CD2 | RPA-2.10 | Mouse IgG1, $\kappa$ | 3.30 |
| CD4 | OKT4 | Mouse IgG2b, $\kappa$ | 1.48 |
| CD6 | MEM-98 | Mouse IgG1 | 4.79 |
| CD9 | HI9a | Mouse IgG1, $\kappa$ | 3.12 |
| CD11a (LFA-1) | TS1/22 | Mouse IgG1 | 1.31 & 6.15 |
| CD27 | O323 | Mouse IgG1, $\kappa$ | 3.70 |
| CD28 | CD28.2 | Mouse IgG1, $\kappa$ | 1.00 |
| CD38 | HIT2 | Mouse IgG1, $\kappa$ | 4.00 |
| CD45 | HI30 | Mouse IgG1, $\kappa$ | 4.50 |
| CD49d | 9F10 | Mouse IgG1, $\kappa$ | 3.25 |
| CD63 (LAMP-3) | H5C6 | Mouse IgG1, $\kappa$ | 3.70 |
| CD70 | 113-16 | Mouse IgG1, $\kappa$ | 3.46 |
| CD80 | 2D10 | Mouse IgG1, $\kappa$ | 1.29 & 4.00 |
| CD81 (TAPA-1) | 5A6 | Mouse IgG1, $\kappa$ | 4.15 |
| CD82 | ASL-24 | Mouse IgG1, $\kappa$ | 4.20 |
| CD107a (LAMP-1) | H4A3 | Mouse IgG1, $\kappa$ | 4.00 |
| CD134 (OX40) | Ber-ACT35 (ACT35) | Mouse IgG1, $\kappa$ | 5.00 |
| CD154 (CD40L) | 24-31 | Mouse IgG1, $\kappa$ | 3.80 |
| CD278 (ICOS) | C398.4A | Armenian Hamster IgG | 3.10 |
| CD317 (BST2) | RS38E | Mouse IgG1, $\kappa$ | 4.10 |
| CD357 (GITR) | 621 | Mouse IgG1, $\kappa$ | 3.60 |
| HLA-A,B,C | W6.32 | Mouse IgG2a, $\kappa$ | 4.50 |

**Table S2.** Surface CD40L levels and densities on T cells exposed to different BSLB substrates (either No UCHT1-Fab ± CD40 or UCHT1-Fab ± CD40).

| | Surface CD40L molec. on CD4<sup>+</sup> T cell | | | | CD40L density on CD4<sup>+</sup> T cells (molec./$\mu\text{m}^2$)* | | | |
| --- | --- | --- | --- | --- | --- | --- | --- | --- |
| | No UCHT1-Fab | | UCHT1-Fab 400 molec./$\mu\text{m}^2$ | | No UCHT1-Fab | | UCHT1-Fab 400 molec./$\mu\text{m}^2$ | |
|  | CD40-BSLB | CD40+BSLB | CD40-BSLB | CD40+BSLB | CD40-BSLB | CD40+BSLB | CD40-BSLB | CD40+BSLB |
| **Average** | 785.7 | 441.8 | 1298.8 | 612.6 | 2.4 | 1.3 | 3.9 | 1.8 |
| **SD** | 459.7 | 286.9 | 595.8 | 422.4 | 1.4 | 0.9 | 1.8 | 1.3 |

**Table S3.** Estimated number of CD40L molecules per SE and estimated CD40L densities on SE using parameters obtained by dSTORM and quantitative FCM.

| | CD40L molec./SE* (±SD) | | CD40L density on patches (molec./$\mu\text{m}^2$)** (±SD) | |
| --- | --- | --- | --- | --- |
| | UCHT1-Fab density (molec./$\mu\text{m}^2$) | | UCHT1-Fab density (molec./$\mu\text{m}^2$) | |
| BSLB composition | 400 | 1800 | 400 | 1800 |
| ICAM-1 | 0.7 ± 0.5 | 2.3 ± 1.6 | 32.2 ± 21.4 | 103.9 ± 73.3 |
| ICAM-1 +CD40 +ICOSL | 12.2 ± 4.3 | 17.6 ± 4.4 | 551.6 ± 196.1 | 793.4 ± 200.4 |
| ICAM-1 +CD40 | 11.1 ± 5.8 | 17.4 ± 9.0 | 499.3 ± 262.4 | 783.9 ± 407.5 |

**Table S4.** Size distribution of SUVs as determined by Nanoparticle Tracking Analyses using light scattering and Brownian motion.

|  | Nanoparticle Tracking Analyses |  |
| --- | --- | --- |
|  | Mean (nm) | SD (nm) |
| Small Unilamellar Vesicles (SUVs/Liposomes) | 85.6 | 25.25 |

